# Patient Similarity Network of Multiple Myeloma Identifies Patient Sub-groups with Distinct Genetic and Clinical Features

**DOI:** 10.1101/2020.06.02.129767

**Authors:** Sherry Bhalla, David T. Melnekoff, Jonathan Keats, Kenan Onel, Deepu Madduri, Joshua Richter, Shambavi Richard, Ajai Chari, Hearn Jay Cho, Joel T. Dudley, Sundar Jagannath, Alessandro Laganà, Samir Parekh

**Affiliations:** Tisch Cancer Institute, Icahn School of Medicine at Mount Sinai, New York, NY, USA; Department of Genetics and Genomic Sciences, Icahn School of Medicine at Mount Sinai, New York, NY, USA; Institute for Data Science and Genomic Technologies, Icahn School of Medicine at Mount Sinai, New York, NY, USA; Translational Genomics Research Institute (TGen), Phoenix, AZ, USA; Department of Hematology and Medical Oncology, Icahn School of Medicine at Mount Sinai, New York, NY, USA; Department of Pediatric Hematology and Oncology, Icahn School of Medicine at Mount Sinai, New York, NY, USA; Department of Pathology, Molecular and Cell Based Medicine, Icahn School of Medicine at Mount Sinai, New York, NY, USA; Tempus Labs, Inc., Chicago, IL, USA; Department of Oncological Sciences, Icahn School of Medicine at Mount Sinai, New York, New York, USA

## Abstract

The remarkable genetic heterogeneity of Multiple Myeloma (MM) poses a significant challenge for proper prognostication and clinical management of patients. Accurate dissection of the genetic and molecular landscape of the disease and the robust identification of homogeneous classes of patients are essential steps to reliable risk stratification and the development of novel precision medicine strategies. Here we introduce MM-PSN, the first multi-omics Patient Similarity Network of newly diagnosed MM. MM-PSN has enabled the identification of three broad patient groups and twelve distinct sub-groups defined by five data types generated from genomic and transcriptomic patient profiling of 655 patients. The MM-PSN classification uncovered novel associations between distinct MM hallmarks with significant prognostic implications and allowed further refinement of risk stratification. Our analysis revealed that gain of 1q is the most important single lesion conferring high risk of relapse, and its association with an MMSET translocation is the most significant determinant of poor outcome. We developed a classifier and validated these results in an independent dataset of 559 pts. Our findings suggest that gain of 1q should be incorporated in routine staging systems and risk assessment tools. The MM-PSN classifier is available as a free resource to allow for an easy implementation in most clinical settings.

## INTRODUCTION

Multiple Myeloma (MM) is an incurable malignancy of bone marrow terminally differentiated plasma cells, affecting more than 30,000 patients each year in the United States, with a median survival of approximately 6 years^1,2^. While the majority of patients initially respond to standard of care therapy, most relapse and become refractory to treatment as they undergo multiple lines of therapy^3^. In particular, about 15% of patients fall in the *high-risk* category and typically relapse within two years from diagnosis^4^. MM is characterized by remarkable clinical and genomic heterogeneity^5^. Chromosomal translocations involving the immunoglobulin heavy chain (IgH) gene locus, hyperdiploidy and complex structural alterations are hallmarks of MM with significant prognostic impact. Recent studies based on next-generation sequencing have revealed more complex patterns of genetic alterations across patients^6,7^, and novel precision medicine approaches, where treatment is guided by the genomic profile of the individual patient, hold great promise for MM therapy^8^. Accurate classification of MM patients into biologically homogeneous classes is thus essential for diagnosis, prognosis and clinical management.

Several classifications of MM based on gene expression have been proposed in the past two decades, such as the TC classification based on cyclin D genes and IgH translocations, the UAMS (University of Arkansas for Medical Sciences) classification, which is based on unsupervised hierarchical clustering and comprises 7 molecular sub-groups enriched for clinically relevant features, and a classification proposed by the HOVON study group, which further extends the UAMS classification to 10 sub-groups^9,10,11^.

Our recent co-expression network model of newly diagnosed MM, MMNet, revealed a clear molecular separation between patients with Ig translocations and hyperdiploidy and identified three novel subtypes characterized by cytokine signaling (CK), immune signatures (IMM) and MYC translocations (MYC)^12^. Another recent study investigated novel MM subtypes based on a targeted DNA panel and revealed a large cluster comprising most HD and IgH translocated patients, and two smaller clusters, one enriched for IgH translocations and one mostly composed of HD patients^13^.

All these findings confirm the significant heterogeneity of MM and provide further evidence for multiple genetic and molecular subtypes of MM. Moreover, the different results obtained from different data types, e.g. DNA vs RNA, suggest that an integrative approach including different omics might further improve patient classification and reveal biologically and clinically informative subtypes of the disease. Recently, Patient Similarity Networks (PSN) have emerged as a powerful tool to capture and structure the complexity and diversity of clinical, genetic and molecular information across a patient population^14^. In a PSN, patients are represented as nodes in a network, much like in a social network, and connected with one another based on how similar their genomic and transcriptomic profiles are. The network structure enables effective identification of communities of highly similar patients, allowing a more comprehensive classification than other approaches based on a single measurement, like gene expression or CNAs. PSNs have been successfully employed to dissect the genomic and molecular complexity of several cancers including medulloblastoma, glioblastoma multiforme, pancreatic ductal adenocarcinoma and metastatic colorectal cancer^15–18^.

Here we describe MM-PSN, a novel PSN of newly diagnosed MM based on multi-omics data from the MMRF CoMMpass study. We identified 3 patient groups and 12 sub-groups, revealing novel insights into co-occurrence of specific genetic lesions and their impact on prognosis and risk stratification.

## METHODS

### Data set acquisition and primary data generation

Whole-Genome Seq (WGS), Whole-Exome Seq (WES) and RNA-Seq data were generated from bone marrow aspirates (CD138+ cells) and peripheral blood (control) of 655 newly diagnosed MM patients enrolled in the MMRF CoMMpass study. Copy Number Alterations (CNA) and translocations were identified from WGS data; somatic Single Nucleotide Variations (SNV) were identified from WES data; gene-level counts and gene fusions were inferred from RNA-Seq data. Sequencing data were integrated with clinical annotation (see Supplementary Methods for details).

### Generation and clustering of Patient Similarity Network

For the generation of MM-PSN we applied the Similar Network Fusion (SNF) method as it did not require *a priori* feature selection and was shown to outperform methods based on single data types as well as other multi-omics approaches such as iCluster^17,19^. We additionally tested the method iClusterBayes^20^ but discarded it because it required feature selection, which significantly biases the results. The R package ‘SNFtool’ (v2.3.0) implementing the SNF method was run on 655 MM tumor samples using gene expression (50,495 genes), SNV (57,736 mutations), gene fusion (13,682 fusions), focal CNA (93 features) and Broad CNA data (39 features) (**Fig. 1A**). Spectral clustering was applied to the fused matrix generated by SNF to determine patient groups, then re-applied to the fused matrices of each group to determine the sub-groups.

**Fig. 1.**
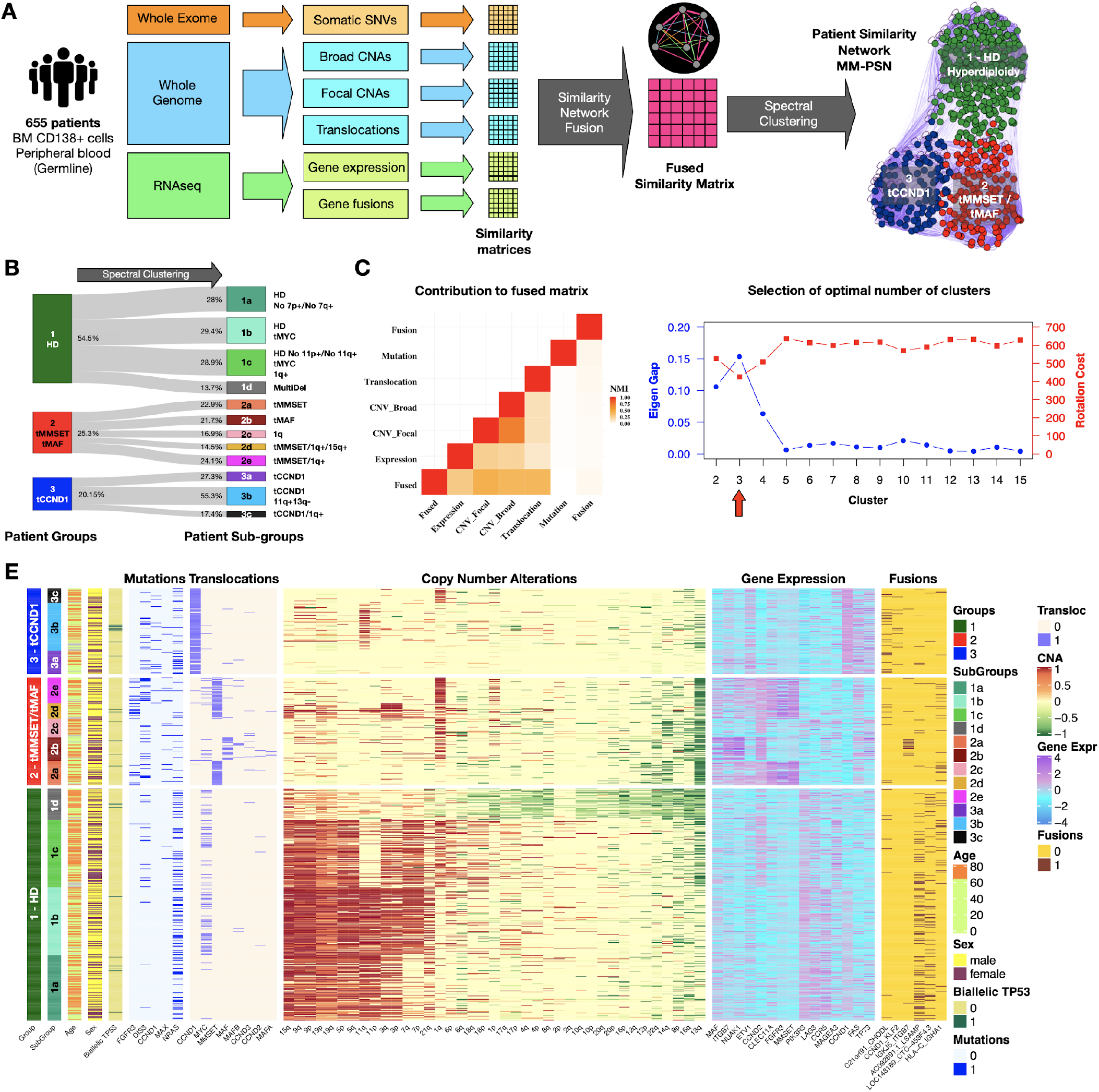
Network generation and identification of groups and sub-groups. **A.** Somatic genetic variants and transcriptomic features from Whole Exome (WES), Whole Genome (WGS) and RNAseq data from 655 patients in the MMRF CoMMpass study was used to generate a Patient Similarity Network (MM-PSN) using the SImilarity Network Fusion approach. Edges connecting patients in the network represent similarity based on one or more feature type (e.g. orange edges in the sample network represent similarity based on SNVs, magenta edges represent similarity based on all the types of features). Spectral clustering was employed to identify patient groups and then re-applied to identify sub-groups enriched for specific features. **B**. Representation of MM-PSN where nodes (patients) are colored according to the three main groups identified by spectral clustering. **C**. The plot shows contribution of the different data types to the fused matrix, in terms of Normalized Mutual Information (NMI). **D**. Eigen Gap (max) and Rotation Cost (min) were used to determine 3 as the optimal number of clusters. **E**. Overview of MM-PSN patient groups and sub-groups. The heatmap shows characterization of the three main groups and twelve sub-groups of MM-PSN based on their enrichment for the different genomic and transcriptomic features.

### Statistical analysis

Progression-free survival (PFS) and overall survival (OS) were analyzed by the Kaplan-Meier method and p-values were calculated using the log-rank test. The Hazard Ratio (HR) was calculated using the Cox proportional hazards method (coxph function from the R package ‘survival’). Associations between categorical covariates and groups/sub-groups were assessed by the Fisher’s exact test. Correlation with continuous variables was assessed using non-parametric statistical measures like the Wilcoxon test (for two variables) or Kruskal-Wallis test (for multiple variables). All statistical analyses were implemented in the R environment.

### Differential feature analysis and functional enrichment

We calculated the differentially expressed (DE) genes for the 12 subgroups by comparing gene expression in each sub-group with the average expression in the other 11 subgroups using the R package ‘edgeR’^21^. Pathway enrichment analysis was performed using the tool ‘g:Profiler’ and visualized as Enrichment Map in Cytoscape^22,23,24^. Pathway definitions were retrieved from Reactome^25^. The CERES scores estimating gene-dependency levels from CRISPR-Cas9 essentiality screens were retrieved from DepMap. Genes with CERES score < −0.5 were considered essential^26,27^.

### MM-PSN classifiers

The dataset was divided into training and validation sets in the ratio of 70:30. The classifier was built by combining three types of features, i.e. focal CNA, translocations and gene expression. Feature selection on copy number values was performed using the Recursive Feature Elimination (RFE) strategy. The top 50 CNA features were selected to maximize the performances. For gene expression, we retained 109 features selected by a two-step feature selection using a Support Vector Classifier with L1 norm (SVC-L1) and RFE. All translocations were retained. We used a stacking classifier with Random forest, Xgboost and linear SVC as base learners and a linear SVM as final estimator.

To validate MM-PSN on an independent gene expression dataset, we trained classification models on the 655 samples using gene expression data only using Logistic Regression, Random Forest and Support Vector Machines (SVM) and compared their performances. The best models were selected using 10-fold cross validation at the group level and 5-fold cross validation at the sub-group level.

### Data and code availability

Sequencing data are available on dbGaP (http://www.ncbi.nlm.nih.gov/gap) under accession number phs000748. The MM-PSN classifier is freely available at https://github.com/laganalab/MM-PSN. The dataset used for validation was retrieved from NCBI GEO (GSE2658).

## RESULTS

### Multi-omics patient similarity network of newly diagnosed MM identifies 3 main patient groups and 12 sub-groups associated with distinct molecular and clinical features

We generated MM-PSN, a multi-omics Patient Similarity Network based on Whole-Exome Seq (WES), Whole-Genome Seq (WGS) and RNA-Seq data from 655 tumor samples from newly diagnosed MM patients enrolled in the MMRF CoMMpass study, using the similarity network fusion (SNF) method^17^ (see **Table 1** for summary patient characteristics). In MM-PSN, each node represents a patient and connecting edges represent similarity based on multiple data types. In particular, for each sample we used: 1) gene expression and 2) gene fusion data from RNA-Seq; 3) somatic single nucleotide variations (SNVs) from WES, 4) Copy Number Alterations (CNAs; focal and broad) and 5) translocations from WGS (**Fig. 1A, 1B**). Translocations and CNAs provided the strongest contribution to MM-PSN, followed by gene expression, gene fusions and SNVs (**Fig. 1C**). We then applied spectral clustering to determine groups of highly similar patients sharing features across the five data types. Our evaluation of the network using Eigen Gap and Rotation Cost suggested three as the optimal number of clusters (**Fig. 1D**; see Methods and Supplementary Methods for details). The three clusters were enriched for (1) Hyperdiploidy (HD) and the t(8;14) translocation of *MYC* (tMYC); (2) translocations t(4;14) of *MMSET* (tMMSET) and t(14;16) of MAF (tMAF); and (3) translocation t(11;14) of *CCND1* (tCCND1), respectively (**Fig. 1A, 1E**). We labeled each group based on these features. Group 1 (HD) included n=357 patients (54.5%) and was enriched for mutations in *NRAS* and gene fusions involving *LSAMP* and *RPL18*; Group 2 (tMMSET+tMAF) included n=166 patients (25.3%) and was enriched for gain(1q) and mutations in *FGFR3*, *DIS3* and *MAX*; Group 3 (tCCND1) included n=132 patients (20.15%) and was enriched for mutations in *CCND1* and *NRAS*. To further investigate intra-group heterogeneity, we re-applied spectral clustering within each group, determining a total of 12 sub-groups (**Fig.1B, 1E**, **Table 2**).

**Table 1.** Patient cohort description and demographics

| Characteristic | Value |
| --- | --- |
| **No. of Patients** | 655 |
| **Age** | 63 (31-93) |
| **Gender** |  |
| **M** | 382 (50.1%) |
| **F** | 218 (33.3%) |
| **N/A** | 109 (16.6%) |
| **Ethnicity** |  |
| **White** | 420 (64.1%) |
| **Black or African American** | 83 (12.7%) |
| **Asian** | 12 (1.8%) |
| **Filipino** | 1 (0.2%) |
| **Honduras** | 1 (0.2%) |
| **Middle Eastern** | 1 (0.2%) |
| **N/A** | 137 (20.9%) |
| **Disease Stage (ISS)** |  |
| **I** | 227 (34.7%) |
| **II** | 226 (34.5%) |
| **III** | 182 (27.8%) |
| **N/A** | 20 (3.1%) |
| **Disease Stage (rISS)** |  |
| **I** | 128 (19.5%) |
| **II** | 339 (51.8%) |
| **III** | 63 (9.6%) |
| **N/A** | 125 (19.1%) |
| **Translocations** |  |
| **MMSET** | 88 (13.4%) |
| **CCND3** | 9 (1.4%) |
| **MYC** | 97(14.8%) |
| **MAFA** | 4 (0.6%) |
| **CCND1** | 126(19.2%) |
| **CCND2** | 6 (0.9%) |
| **MAF** | 27 (4.1%) |
| **MAFB** | 10 (1.5%) |
| **None** | 307 (46.9%) |
| **Multiple \*** | 19 (2.9%) |
| **Disease Status** |  |
| **PFS < 1yr** | 225 (34.4%) |
| **PFS < 2yr** | 365 (55.7%) |

\*CCND1 + MYC: 5; CCND2 + MYC: 2; MAF + MYC: 5; MAFB + MYC: 2; MMSET + MAF: 1; MMSET + MYC: 4

**Table 2.**
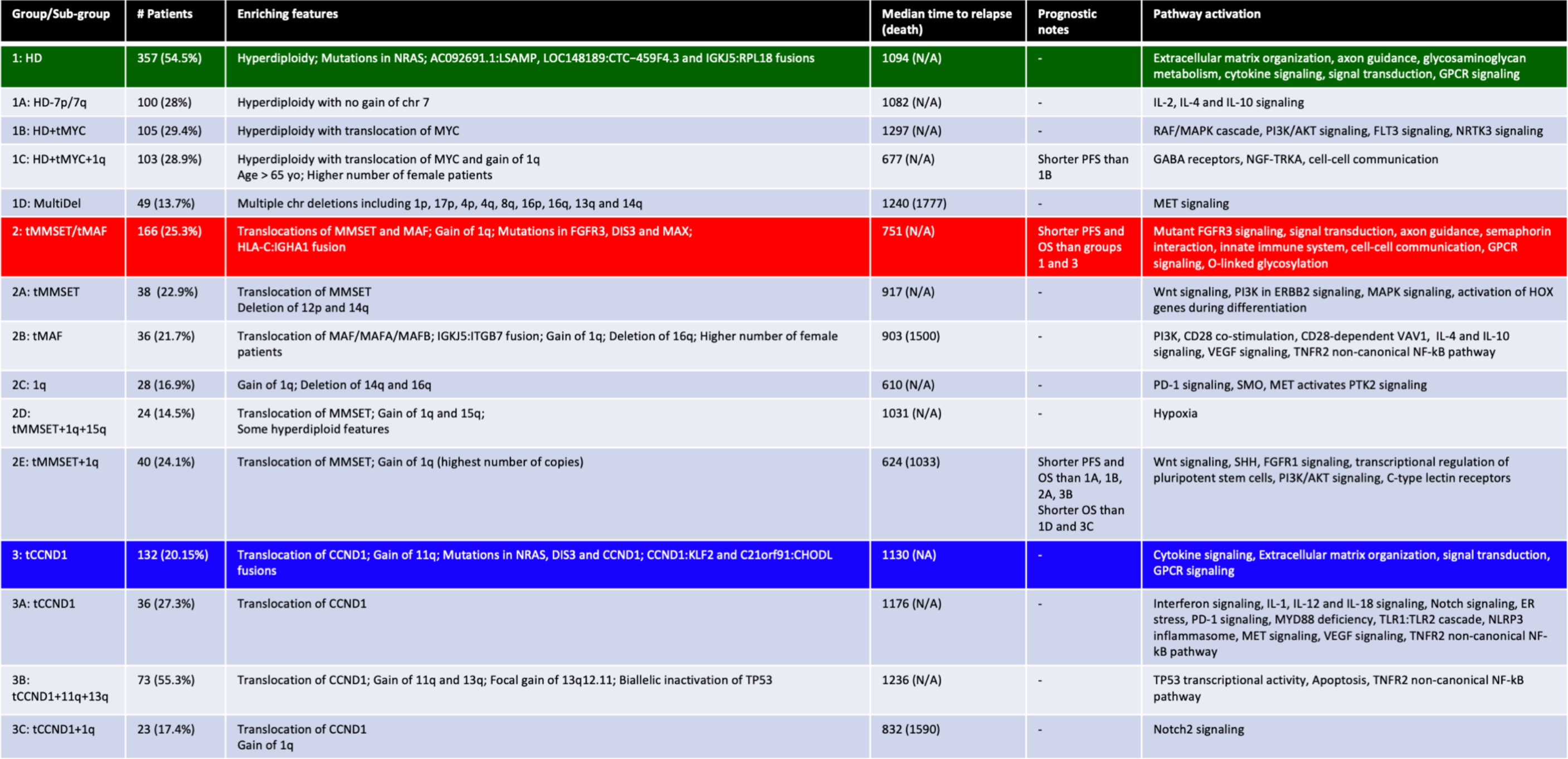
Summary of the MM-PSN sub-groups.

Group 1 was comprised of four sub-groups. Sub-groups 1a (n=100; 28%), 1b (n=105; 29.4%) and 1c (n=103; 28.9%) were mostly characterized by HD with differences in gain of chromosomes 7 and 11, which were almost not detected in sub-groups 1a and 1c, respectively. Sub-groups 1b, c and d were enriched for tMYC, which was virtually absent in sub-group 1a. Additionally, sub-group 1c was significantly enriched for the fusion HLA-C:IGHA1, gain(1q) and del(13q). Sub-group 1d (n=49; 13.7%) had very weak HD signal and was significantly enriched for multiple chromosome deletions instead, including del(1p), del(17p), del(14q), del(16q) and del(13q). Group 2 was comprised of five sub-groups. Sub-groups 2a (n=38; 22.9%), 2d (n=24; 14.5%) and 2e (n=40; 24.1%) were enriched for the tMMSET, while sub-group 2b (n=36; 21.7%) was enriched for translocation of tMAF, tMAF-A and tMAF-B. Sub-group 2b was enriched for the IGKJ5:ITGB7 gene fusion. Del(13q) was observed in all 5 sub-groups, while gain(1q) was significantly enriched in sub-groups 2b, 2c (n=28; 16.9%), 2d and 2e. Group 2d was also enriched for gain of 15q and showed other weak HD features, such as gain of chromosomes 3 and 19.

Group 3 was comprised of three sub-groups, all enriched for tCCND1. Sub-group 3a (n=36; 27.3%) had virtually no CNAs, while sub-group 3b (n=73; 55.3%) was additionally enriched for del(13q), specifically focused at 13q12.11, TP53 biallelic inactivation (mutation + deletion), gain(11q) and other sparse CNAs. Sub-group 3c (n=23; 17.4%) was enriched for gain(1q).

We compared MM-PSN with the UAMS and our previous MMNet classifications^10,12^ (Supp Fig. 1A, 1B). Although the results showed significant agreement among the three classification systems, MM-PSN revealed a more granular view of patient classes and their defining features.

### Survival analysis of MM-PSN identifies co-occurrence of genetic lesions and 1q gain with prognostic impact

Survival analysis of the three Groups showed poorer PFS in patients from Group 2(tMMSET+tMAF), compared to the other two groups (**Fig. 2A, 2B**). Analysis performed within each group revealed significant differences in survival among sub-groups. In particular, gain(1q) identified a sub-group of HD patients, 1c(HD+tMYC+1q), with significantly shorter PFS within group 1 (HR=1.6; p=0.04), and significantly shorter OS than patients in sub-group 1b(HD+tMYC) (HR=2.26, p=0.01) (**Fig. 2C, 2D**). Within group 2(tMMSET+tMAF), patients in sub-group 2e(tMMSET+1q) had significantly worse PFS (HR=2.04, p=0.002) and OS (HR=2.71, p=5e-04) (**Fig. 2E, 2F**). No significant differences in survival were observed among sub-groups of Group 3, although sub-group 3c(tCCND1+1q) had a slightly poorer PFS (HR=1.3, p=0.3) (**Fig. 2G, 2H**). Overall, sub-group 2e had the poorest outcome both in terms of PFS and OS (**Fig. 2I, 2J**).

**Fig. 2.**
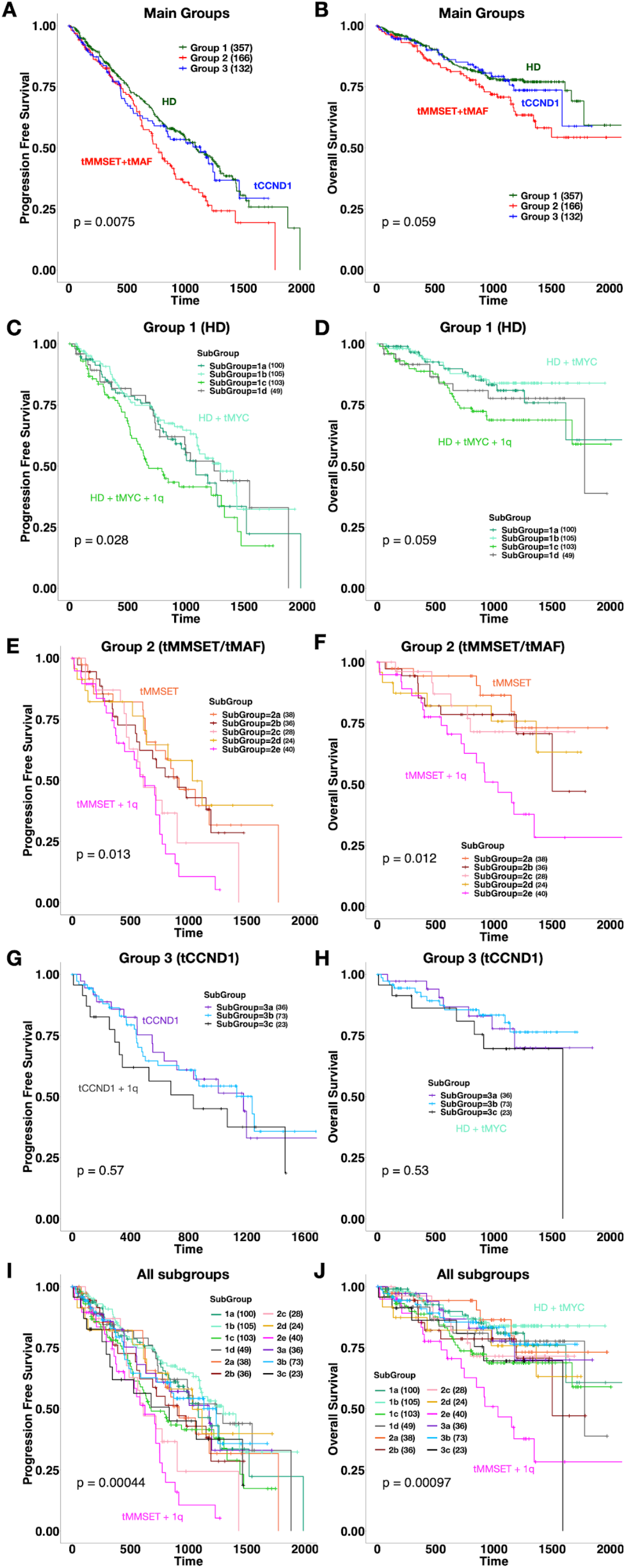
Survival analysis identifies sub-groups enriched for 1q gain associated with poor outcomes. **A, B.** Progression Free Survival (PFS) and Overall Survival (OS) plots for the three main patient groups identified by MM-PSN, showing significant poorer outcome for the tMMSET+tMAF group. **C, D**. Survival plots for sub-groups of group 1 show shorter PFS in patients from sub-group 1C characterized by hyperdiploidy, tMYC and 1q gain, compared to patients in sub-group 1B which do not have 1q gain. **E, F**. Survival plots for sub-groups of group 2 show shorter PFS and OS in patients from sub-group 2E, enriched for tMMSET and 1q gain. **G, H**. Survival plots for sub-groups of group 3 do not show significant differences in either PFS or OS. **I, J**. Survival plots for all twelve sub-groups of MM-PSN, indicating poorer outcome of patients in sub-group 2E.

### The gain of 1q is the most significant independent determinant of poor outcome

Out of 45 patients with both tMMSET and gain(1q), 40 (89%) were in sub-group 2e. We found that, overall, patients with both tMMSET and gain(1q) (n=45) had much shorter PFS and OS compared to patients with tMMSET alone (n=43) (PFS: HR=2.3, p=0.005; OS: HR=3.47, p=0.008) (**Fig. 3A, 3B**). Since the number of 1q copies has been previously reported as being prognostically relevant^28,29^, we evaluated it across all the sub-groups. Sub-group 2e had the highest number of copies (p=0.0001) (Supp Fig. 2A). Stratification of all patients in CoMMpass with CNA data available (n=870), confirmed that patients carrying 4 copies of 1q had worse PFS and OS than patients carrying 3 copies (PFS: HR=1.4, p=0.06; OS: HR=1.6, p=0.04,) (Supp Fig. 2B, 2C). Next, considering the poorer prognosis observed in HD patients with gain(1q) in sub-group 1c, we evaluated the association of HD and 1q across the whole cohort. The analysis revealed significantly shorter PFS and OS in HD patients with gain(1q), as compared to HD patients with no 1q alterations (PFS: HR=1.56, p=0.008; OS: HR=2.04, p=0.004) (**Fig. 3C, 3D**). These findings indicate a significant role of gain(1q) in driving prognosis. Multivariate cox regression analysis including basic demographics and major alterations enriched in the network, showed that gain(1q) alone and co-occurring with tMMSET were both significantly associated with worse PFS and OS (PFS: HR=2.08, p= <0.001,OS: HR=3.49, p= <0.001) (Supp Figures 3 and 4). Biallelic inactivation of TP53 (mutation and deletion) was also significantly associated with worse prognosis (PFS: HR= 2.02, p=0.005; OS: HR= 3.08, p=0.0002) (Supp Figures 3,4 and 6). The translocation of *MAF*, which has been considered a high-risk alteration in previous studies^30,31^, was not significantly associated with worse outcome. The analysis also showed that African American and male patients had also worse OS (AA: HR=1.73, p=0.012; male: HR=1.50, p=0.041). In contrast, gain(15q), which was a widespread alteration in main group 1 (HD) and significantly enriched in sub-group 2d(tMMSET+1q+15q), was associated with better PFS and OS (PFS: HR=0.66, p=3e-04; OS: HR=0.7, p=0.04) (**Fig. 3E, 3F**). This might in part explain the better prognosis observed in sub-group 2d compared to 2e, despite the presence of tMMSET and 1q gain.

**Fig. 3.**
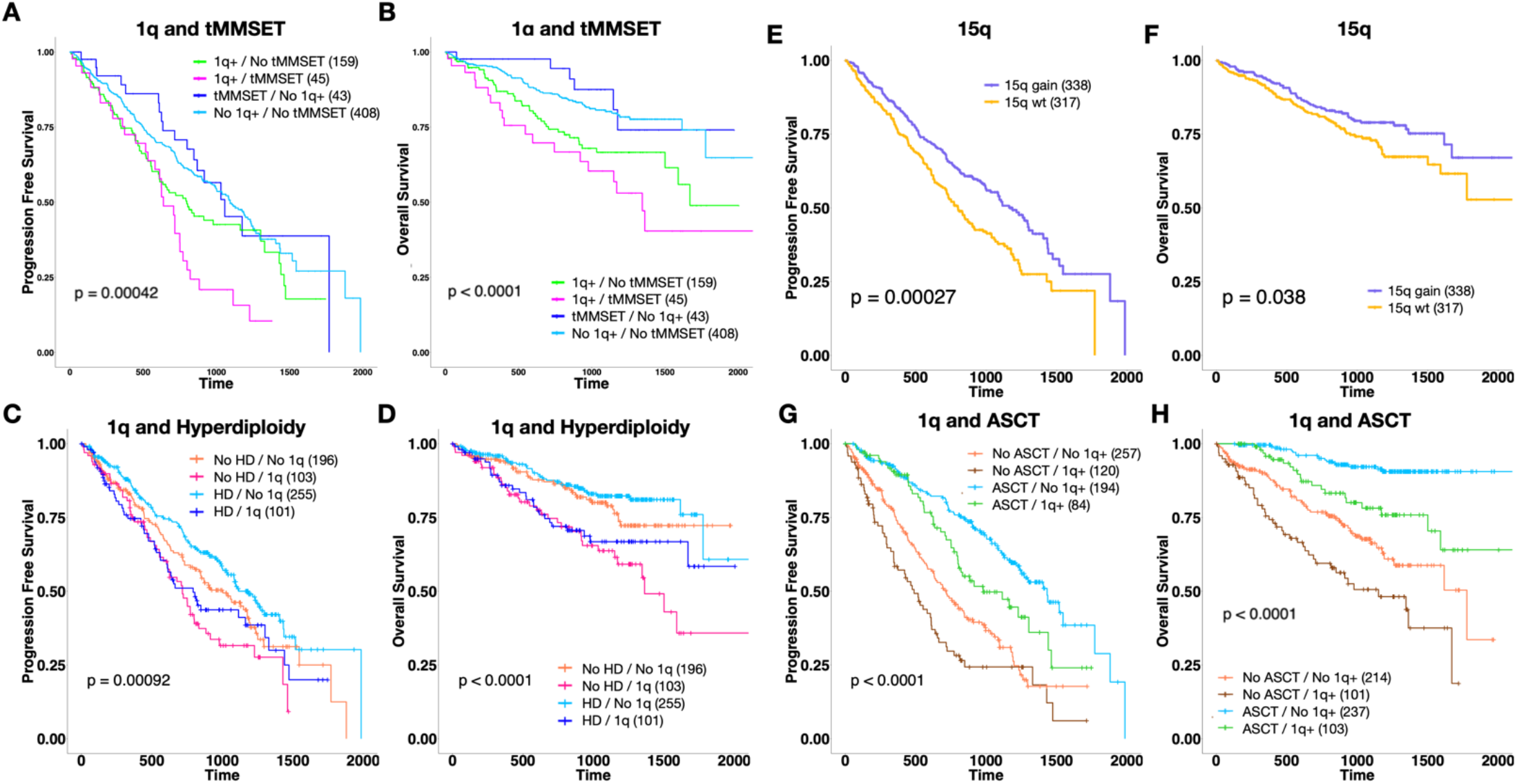
Prognostic implications of 1q, tMMSET and 15q. **A, B.** survival plots show that patients with 1q gain with or without tMMSET have poorer outcome than patients with tMMSET alone. **C, D**. Survival plots show 1q gain identifies a subset of hyperdiploid patients with significantly shorter PFS and OS. **E, F**. Gain of 15q is associated with better PFS and OS. **G, H**. Survival plots show that patients with 1q gain that received ASCT have significantly better PFS and OS compared to patients that didn’t receive ASCT.

### The negative prognostic impact conferred by 1q gain is not overcome by treatment

We asked whether treatment influenced outcome for patients with gain(1q) through multivariate cox regression. Neither induction therapy (e.g. bortezomib/carfilzomib with lenalidomide) nor autologous stem cell transplant (ASCT) could overcome the negative prognostic impact of gain(1q). While patients with gain(1q) who received ASCT (n=108) had better outcomes compared to patients with gain(1q) who did not receive it (n=217), the prognosis was still significantly poorer than in patients without gain(1q) who received ASCT (n=194) (**Fig. 3G, 3H**, Supp Figures 3, 4). Across the whole cohort (n = 655), we observed better PFS in patients treated with carfilzomib-based therapies (HR=0.46, p=0.05) and ASCT was significantly associated with both better PFS and OS (PFS: HR=0.26, p=0.001; OS: HR=0.26, p=0.001) (Supp Figures 3 and 4).

### Gain of 1q stratifies risk beyond ISS classes

The International Staging System (ISS) is the most used risk score in the clinical setting and is based on serum beta-2 microglobulin and albumin^32^. The revised ISS (rISS) additionally includes lactate dehydrogenase (LDH) and the high risk cytogenetic markers tMMSET, tMAF and del(17p), but not gain(1q)^31^. Stratification of patients based on gain(1q) and the ISS/rISS classes revealed that gain(1q) could identify patients with significantly shorter OS in all three ISS classes (ISS I: p=0.013; ISS II: p=0.007; ISS III: p=0.02) and in rISS classes II and III (rISS II: p=0.01; rISS III: p=0.002) (**Fig. 4**). Gain(1q) also identified patients at higher risk of relapse in ISS classes I and III (ISS I: p=0.02; ISS III=p=0.03) and rISS classes II and III (rISS II: p=0.004; rISS III: p=0.019) (**Fig. 4**).

**Fig. 4.**
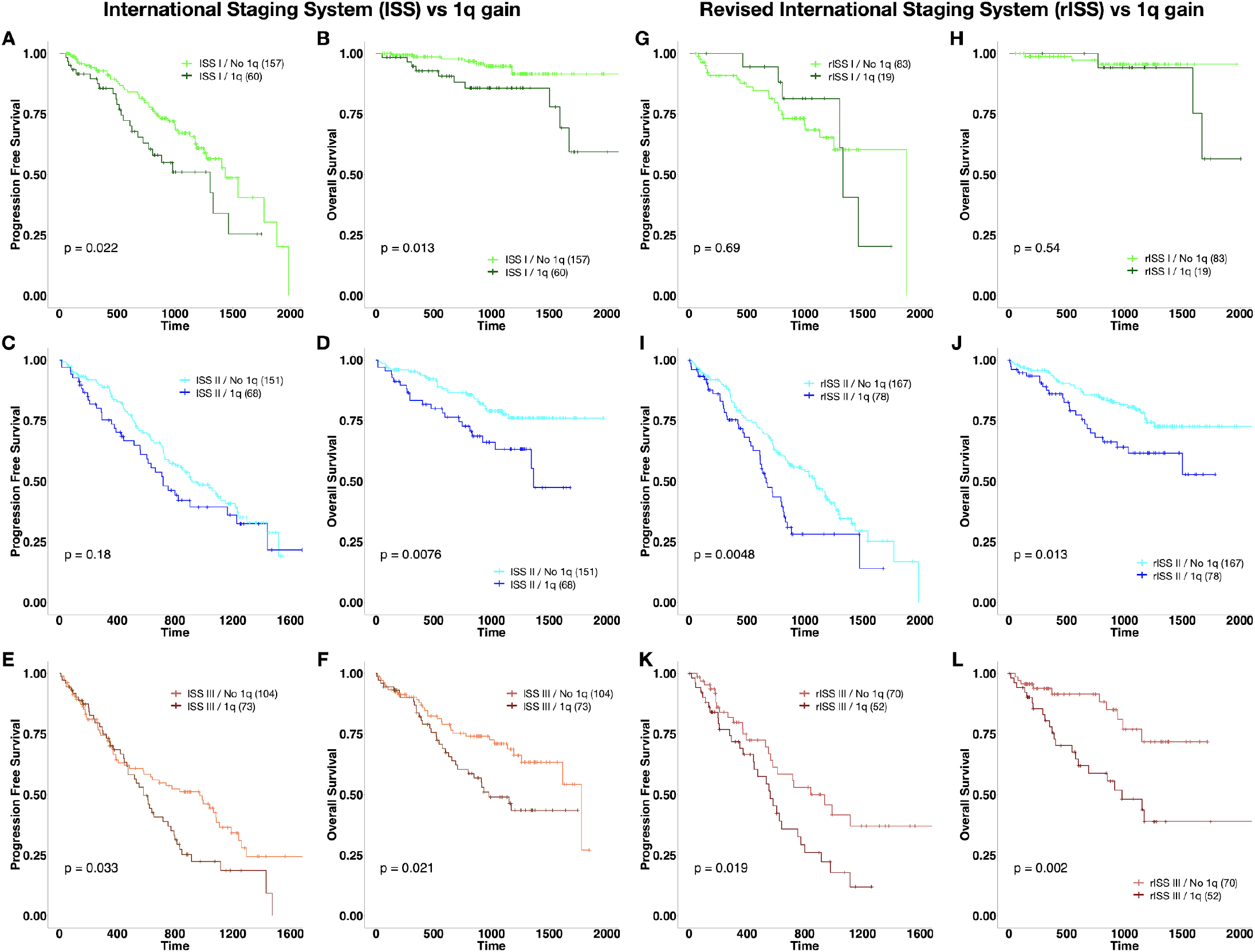
Gain of 1q identifies higher risk patients in lower-risk ISS and rISS classes. **A-F** Gain of 1q significantly stratifies risk for relapse and mortality in patients in ISS classes I and III, and risk of mortality in patients in ISS class II. **G-L**. Gain of 1q xssignificantly stratifies risk for relapse and mortality in patients in rISS class II, and risk of mortality in patients in rISS class III.

### Differential pathway analysis reveals specific enrichment patterns in MM-PSN sub-groups

To characterize the 12 MM-PSN sub-groups functionally we performed pathway activation analysis based on differential gene expression (**Table 2**, **Fig. 5A**, Supp Table 1). The analysis identified 10 main enrichment categories containing most of the activated pathways and revealed convergence of different sub-groups towards broad pathways such as extracellular matrix organization (ECM), signal transduction and cytokine signaling. We observed distinct patterns of pathway activation in sub-groups of Group 1. Sub-group 1a(HD-7p/7q) was enriched for interleukin signaling, including IL-2, IL-4, IL-10 and IL-13, while sub-group 1b(HD+tMYC) was characterized by aberrant activation of PI3K/AKT signaling, which was shared with sub-group 2b(tMAF). Sub-group 1c(HD+tMYC+1q) was enriched for GABA receptor signaling and cell-cell communication, while sub-group 1d(MultiDel) was enriched for MET signaling.

**Fig. 5.**
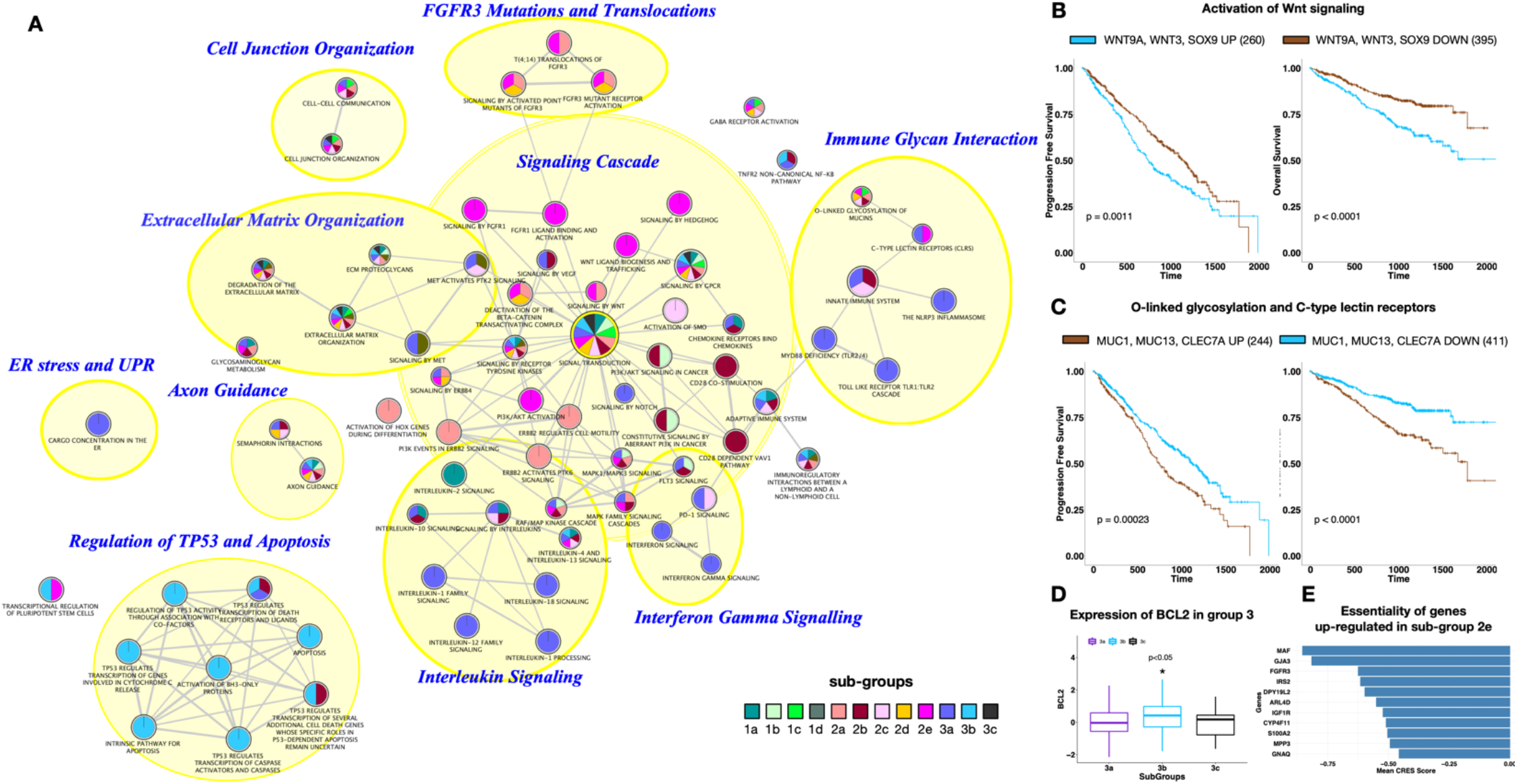
Pathway activation in MM-PSN sub-groups. The figure shows the enrichment map for selected pathways that are significantly activated in MM-PSN sub-groups. Each circle indicates a pathway and the colors represent the sub-groups with significant activation of the pathway. Edges connect pathways that share genes. Most pathways are organized in categories (yellow areas), which are identified by the large blue labels. **B.** Activation of Wnt signaling mediated by the up-regulation of WNT9A, WNT3 and SOX9 observed in sub-group 2e is associated with shorter PFS and OS. **C.** Activation of O-linked glycosylation and C-type lectin receptors mediated by the up-regulation of MUC1, MUC13 and CLEC7A observed in group 2 and sub-group 1c is associated with shorter PFS and OS. **D.** Heterogeneity of BCL2 expression within patients with t(11;14): sub-group 3b had significantly higher expression of BCL2. **E.** Up-regulated genes in sub-group 2e with CERES essentiality score < −0.5.

Group 2 was enriched for growth, proliferation and differentiation pathways, including ERBB2 signaling in sub-group 2a(tMMSET), aberrant FGFR3 signaling in sub-groups 2a(tMMSET), 2d(tMMSET+1q+15q) and 2e(tMMSET+1q), hedgehog signaling in sub-groups 2c(1q) and 2e(tMMSET+1q), and IGF1R and Wnt signaling cascades in sub-group 2e(tMMSET+1q). Multivariate cox regression of Wnt signaling’s components up-regulated in 2e revealed a prognostic impact of *WNT3*, *WNT9A* and *SOX9*. Patients with up-regulation of these three genes had significantly shorter PFS (HR: 1.45; p = 0.001) and OS (HR: 2; p < 0.0001) (**Fig. 5B**). Similarly, up-regulation of genes from the O-linked glycosylation and C-type lectin receptor pathways, *CLEC7A*, *MUC1* and *MUC13*, identified patients with worse PFS (HR: 1.52; p = 0.0002) and OS (HR: 2; p<0.0001) (**Fig. 5C**). Of note, both *MUC1* and *WNT9A* are located in 1q, thus the activation of the glycosylation and Wnt pathways may be driven by gain(1q).

Group 3 was overall enriched for inflammation and immune-related pathways. Specifically, sub-group 3a was mostly characterized by immune and inflammation-related pathways, such as interferon signaling, interleukin signaling including IL-1, IL-10, IL-12 and IL-18, the toll-like receptor cascade and the NLRP3 inflammasome. Sub-group 3c was characterized by Notch2 and MET signaling. Sub-group 3b, instead, was enriched for TP53- and apoptosis-related pathways, in particular the activation of BH3-only protein driven by the up-regulation of *TP63*, *TP73* and *PMAIP1*, which encodes the protein NOXA. Additionally, sub-group 3b had significantly higher expression of *BCL2* within Group 3 (**Fig. 5D**). These findings, along with the enrichment for tCCND1, suggest potential therapeutic implications of the BH3-mimetic BCL2 inhibitor Venetoclax for patients in this sub-group. Finally, since sub-group 2e identified patients with the poorest outcome, we further investigated the essentiality of the genes significantly dysregulated in these patients, as determined by CRISPR-Cas9 screens in MM cell lines with concurrent tMMSET and gain(1q) (OPM2, LP1 and KMS11). We identified essential genes as those with a CERES score below −0.5^26,27^. Among the genes uniquely up-regulated in sub-group 2e, *IGFR1*, which has been previously reported as aberrantly expressed in aggressive MM, had the highest likelihood of being essential (Median CERES score = −0.58)^26,27,33,34^. When we extended the analysis to genes up-regulated in sub-group 2e and any other sub-group in Group 2, we identified 3 additional essential cancer-associated genes: MAF, FGFR3 and GNAQ (Median CERES scores: MAF=−1.16; FGFR3=−0.88; GNAQ=−0.51) (**Fig. 5E**).

### A prognostic gene expression signature induced by MM-PSN predicts survival in an independent dataset

To validate our MM-PSN model, we first trained a gene expression classifier on the 12 MM-PSN sub-groups, employing an approach based on support vector machine (SVM) and ANOVA-based feature selection. Then, we tested the classifier on a gene expression dataset with 559 tumor samples from newly diagnosed patients pre-TT2 and -TT3 treatments. The classifier was able to predict patients in the 3 main groups and to replicate the prognostic findings from MM-PSN, with group 2 having significantly shorter PFS (HR=2.25, p=3e-07) and OS (HR=1.8, p=8e-05) than groups 1 and 3 (Supp Fig. 7A, 7B). Furthermore, we validated sub-group 2e as having significantly shorter PFS (HR=3.08, p=0.004) and OS (HR=3.64, p=0.002) than 2a (Supp Fig. 7C, 7D), supporting the negative prognostic impact of co-occurrence of tMMSET and gain(1q).

## DISCUSSION

Classification of MM patients and risk stratification has relied on either biochemical parameters, cytogenetic markers or gene expression analysis. Recent studies have demonstrated that integrative multi-omics approaches to patient classification and risk stratification can significantly outperform methods based on a single data type^19,35,36^. PSNs have been proposed as a powerful integrative approach to combine different omics data, create a comprehensive view of a given disease and identify disease subtypes ^15,17,37–39^.

In this study we have generated a PSN of newly diagnosed MM patients, called MM-PSN, using five different data types derived from WGS, WES and RNAseq, and have determined a novel classification of MM consisting of 3 main groups and 12 sub-groups.

Our MM-PSN analysis identified several sub-groups enriched for gain(1q), each associated with other recurrent lesions and all of them having the shortest median time to relapse and death compared to the other sub-groups. In particular, co-occurrence of gain(1q) with the t(4;14) MMSET translocation was significantly enriched in sub-group 2e and was the most significant and frequent event associated with poor outcome, as further confirmed in an independent dataset. Gain(1q) is observed in ~40% of MM cases and in several other types of cancer, including breast cancer, hepatocellular carcinoma and myeloproliferative neoplasms, and has been the subject of numerous studies in the past decade^40–43^. A recent study has shown that newly diagnosed MM patients with gain of 1q who received induction therapy with bortezomib, lenalidomide and dexamethasone (VRD) are at higher risk of early relapse, especially when amp(1q) is detected^28^. Another study found that relapsed/refractory MM patients (RRMM) with gain of 1q have worse outcomes after treatment with Daratumumab^44^.

Despite increasing evidence supporting the prognostic relevance of gain(1q), current staging systems such as the International Staging System (ISS) and its revised version, rISS, do not include it^31,32^. Our results show that gain(1q) could significantly stratify patients in almost all risk classes into high vs low risk sub-classes, in terms of both PFS and OS. This is in agreement with a previous study by Walker et al, where the authors showed that integrating copy number and structural abnormalities, including amp(1q), into ISS could further increase its prognostic power^45^. These findings strongly suggest that gain(1q) should be incorporated into these systems and used in the clinic to determine patient risk.

Recent studies such as the Myeloma Genome Project (MGP) as well as the NCRI Myeloma XI and the MRC Myeloma IX trials have described co-occurrence of two high-risk features, termed a ‘double-hit’ event, as a reliable indicator of poor prognosis^29,46^. One such double-hit event is the biallelic inactivation of TP53 through loss and/or mutation. In this study, we identified 23 patients affected by this alteration and survival analysis confirmed the significant poor prognosis in these patients in terms of both PFS and OS. Although these cases were distributed across the three main groups, we observed significant enrichment in the tCCND1 sub-group 3b (n=7; 30%). However, since patients with biallelic TP53 inactivation were only a small fraction of the patients in the sub-group, the presence of the alteration did not have a significant impact on the outcome of the sub-group overall.

Functional characterization of the MM-PSN sub-groups through pathway activation analysis based on differential gene expression revealed meaningful insights with important biological and clinical implications. The enrichment for the O-linked glycosylation of mucins and C-type lectins observed across all of group 2 (tMMSET/tMAF) and sub-group 1c(HD+tMYC+1q) was defined by the dysregulation of distinct genes. This was in agreement with our previous network analysis demonstrating a potential role of the c-type lectin gene *CLEC11A* as an upstream regulator of tMMSET-driven MM^12^. Up-regulation of *MUC1* was a common event between group 2 (tMMSET/tMAF) and sub-group 1c(HD+tMYC+1q), possibly driven by gain(1q), which was significantly enriched in both patient classes. Previous studies have shown aberrant expression of *MUC1* in MM as a driver event causing the activation of the NF-κB pathway, the stabilization of β-catenin and the regulation of *MYC* expression^48,49^. Furthermore, targeting of MUC1 with the MUC1-C inhibitor GO-203, was reported to overcome resistance to lenalidomide in MM cells^50^.

Sub-group 2e was also enriched for IGF1R signaling cascade, mediated by the up-regulation of *IGF1R*, *IRS2*, *FGF9* and *IRS4*. IGF1R (Insulin-like growth factor-1 receptor) is a receptor tyrosine kinase with high affinity for IGF1 (insulin-like growth factor 1). It has been previously reported as over-expressed in MM patients and human cell lines with tMMSET and its activation promotes cell growth and survival^33,34^. Our analysis identified *IGF1R* as the most essential up-regulated gene in sub-group 2e, according to data from CRISPR-Cas9 screening of MM cell lines with concurrent tMMSET and gain(1q). Therefore, it represents a potential vulnerability that could be exploited therapeutically in this specific class of patients.

Our previous work has demonstrated that demethylation of NOXA induced by the DNA methytransferase inhibitor Decitabine (DAC) could overcome resistance to bortezomib in mantle cell lymphoma^51^. More recent studies have shown that down-regulation of NOXA was associated with resistance to the BH3-mimetic BCL2 inhibitor venetoclax in leukemia and lymphoma cell lines^52^. Moreover, pharmacological induction of NOXA by the HDAC inhibitor panobinostat and by 5-Azacytidine was reported to prime DLBCL and AML cells, respectively, to BCL2 inhibitor-induced cell death^53,54^. In MM, tCCND1 and high BCL2 expression are considered markers of sensitivity to venetoclax^55^. The activation of NOXA in the MM-PSN sub-group 3b may identify more specifically the sub-population of tCCND1/high-BCL2 patients that are sensitive to venetoclax. We are currently investigating this hypothesis *in vitro*.

The MM-PSN classification is a valuable and accessible resource that can be employed in most clinical settings. While our classifier can be easily utilized to predict group and sub-group membership of any MM patient, especially in academic centers that perform RNA-Seq, WES and/or array-based comparative genomic hybridization (aCGH), oncologists in smaller practices can classify patients by the co-occurrence of genetic lesions detected by FISH/Cytogenetics. Finally, the classifier can be implemented into a CLIA certified test for wide adoption by the oncology community. This paves the way for future research into drug repurposing approaches aimed at novel therapies tailored to specific patient sub-groups.

## ACKNOWLEDGEMENTS

The authors thank all the patients enrolled in the MMRF CoMMpass study. This work was supported by NIH-NCI (R21-CA209875-01A1; R01-1R01CA244899-01A1), Tisch Cancer Institute (TCI) (NCI Support Grant: P30 CA196521) and the Multiple Myeloma philanthropic fund. This work was also supported in part through the computational resources and staff expertise provided by Scientific Computing at the Icahn School of Medicine at Mount Sinai.

## AUTHORSHIP

Contribution: S.B., A.L. and S.P. conceived and designed the study; S.B. led the study and performed data analysis; D.T.M. performed pathway analysis; J.K. provided sequencing data; D.M., J.R., S.R., A.C., H.J.C., S.J. and S.P. contributed patient samples and clinical data; K.O. and J.T.D. provided scientific expertise; S.B., A.L. and S.P. wrote the manuscript; all authors revised and approved the final version of the manuscript.

## SUPPLEMENTARY FIGURES

**Supp Fig. 1.**
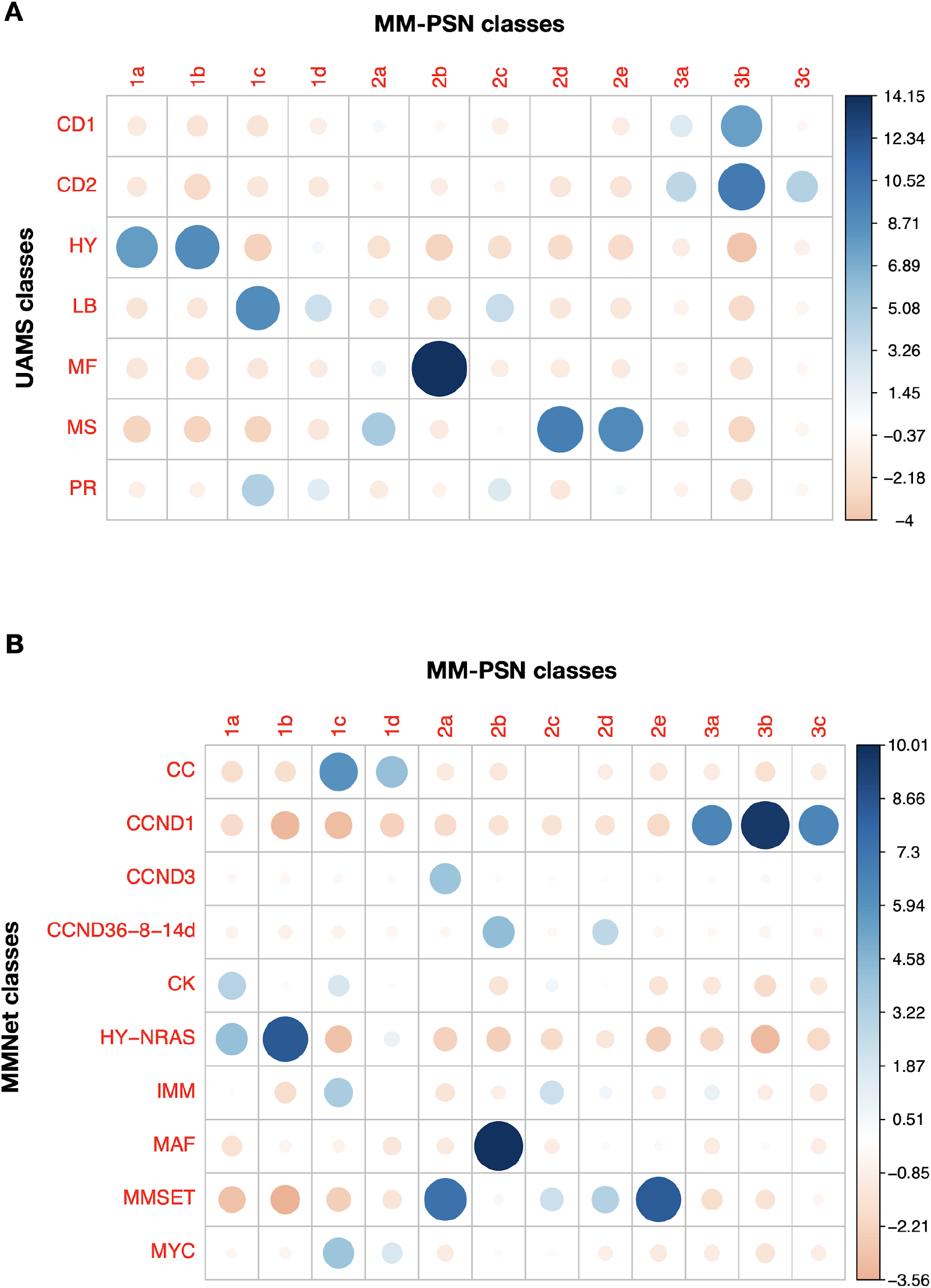
Comparison of MM-PSN sub-groups with UAMS and MMNet classes. The comparison was performed using Pearson residuals. Positive residuals are in blue and specify a positive association between the corresponding classes. Negative residuals are in red and imply a negative association between the corresponding classes. **A.** Comparison between the MM-PSN sub-groups and the UAMS classes (CD1/CD2: CCND1; HY: hyperdiploid; LB: low bone disease; MF: MAF; MS: MMSET; PR: proliferative). **B.** Comparison between the MM-PSN sub-groups and the MMNet classes (CC: cell cycle; CK: cytokines; IMM: immune).

**Supp Fig. 2.**
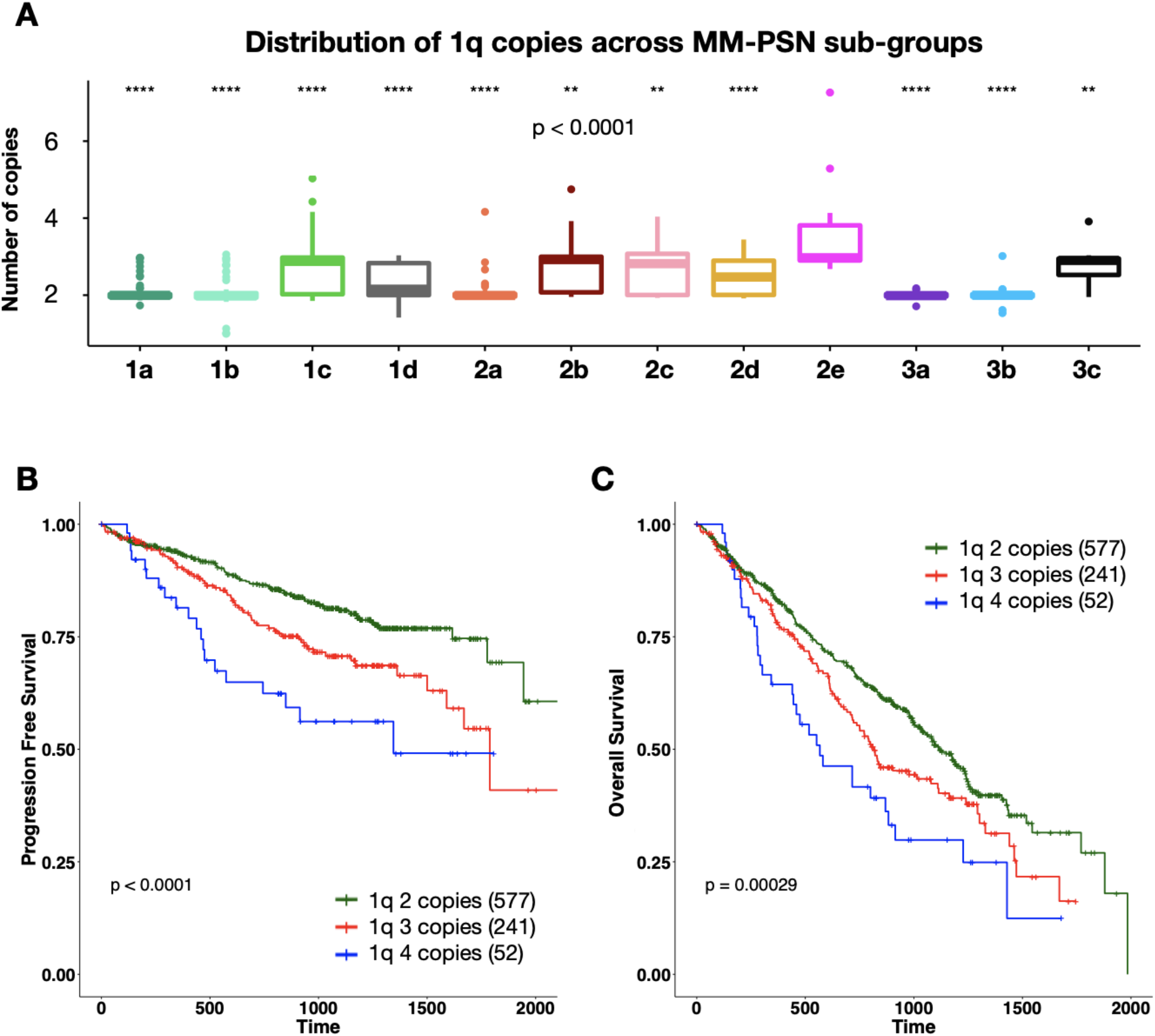
Prognostic implications of 1q gain and amplification. **A**. The distribution of the number of copies of 1q across MM-PSN sub-groups show a significantly higher number of copies in patients in 2e. The stars on top of each bar indicate significance compared to 2e(ns: P > 0.05, *: P ≤ 0.05, **: P ≤ 0.01, ***: P ≤ 0.001, ****, P ≤ 0.0001). **B-C**. Number of 1q copies significantly stratify PFS and OS. Amplification of 1q (4 or more copies) confers much worse prognosis than gain (3 copies).

**Supp Fig. 3.**
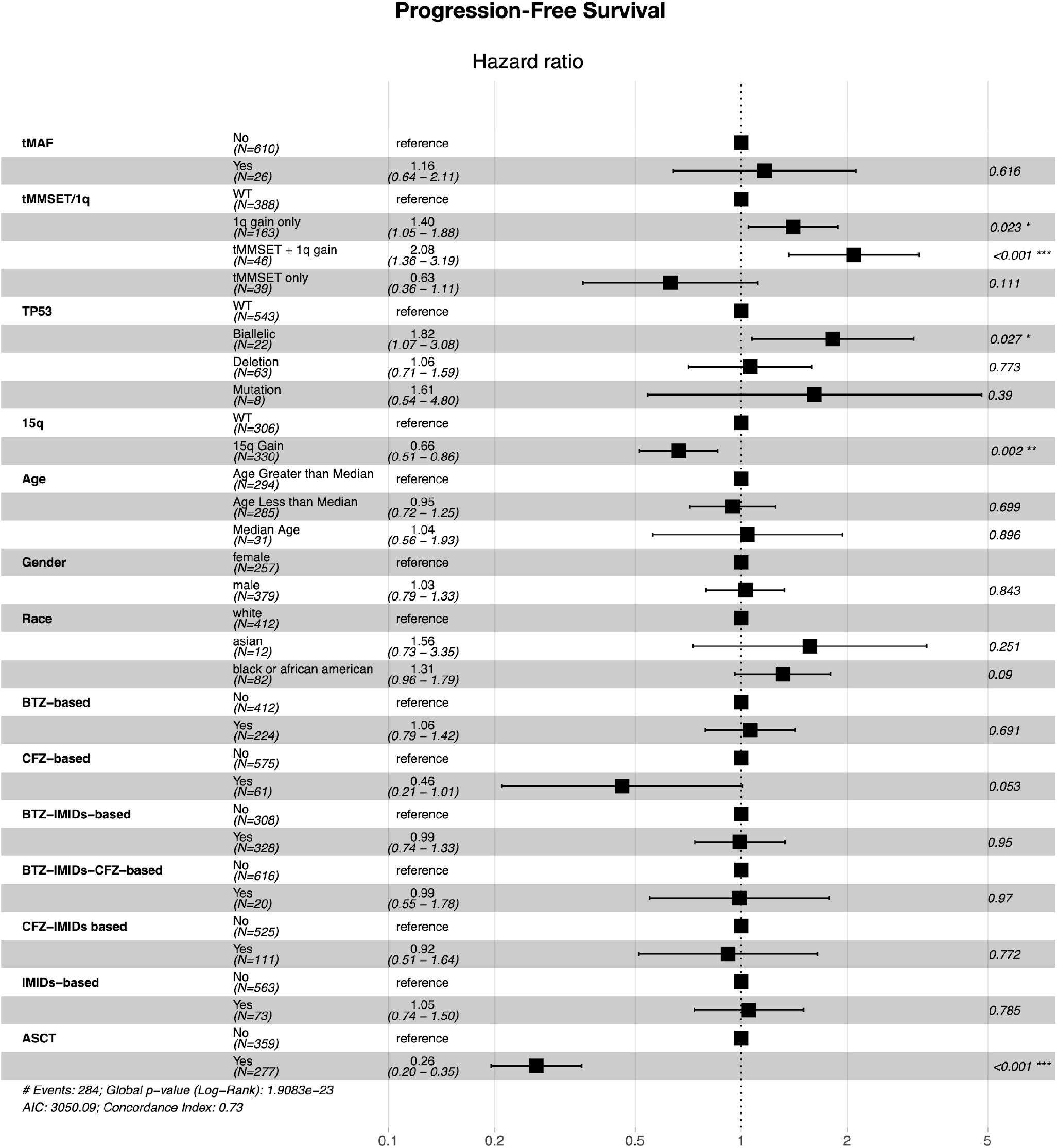
Multivariate cox-regression analysis of progression free survival. The analysis reveals that 1q gain, its combination with tMMSET and biallelic inactivation of TP53 are significantly associated with shorter PFS, while gain of 15q is associated with better PFS. Among treatments, autologous Stem Cell Transplant (ASCT) is significantly associated with better survival in the context of all the high-risk factors included in the model. Carfilzomib-based treatments have borderline significant benefits.

**Supp Fig. 4.**
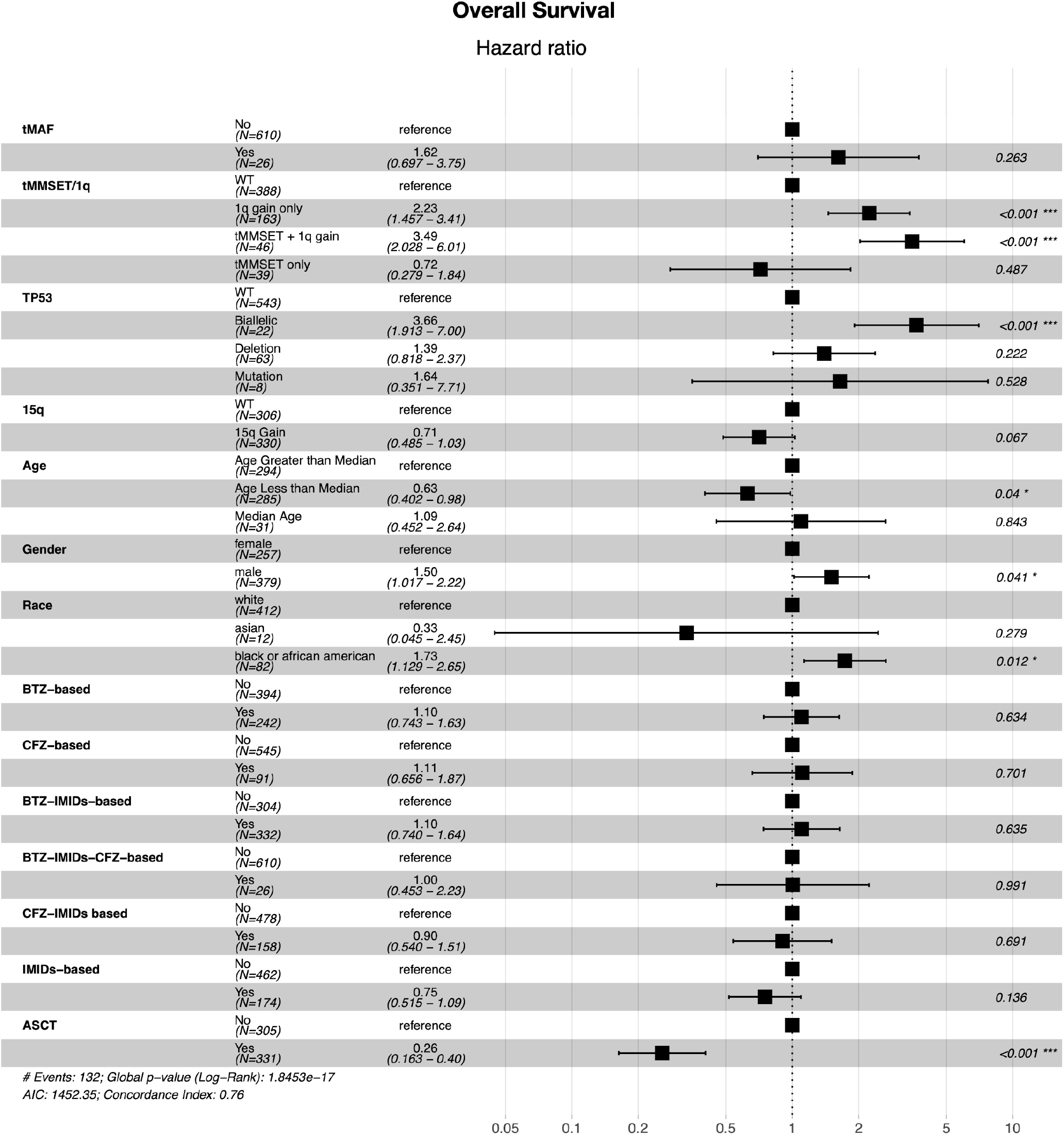
Multivariate cox-regression analysis of overall survival. The analysis reveals that 1q gain, its combination with tMMSET and biallelic inactivation of TP53 are significantly associated with shorter OS. Gain of 15q has borderline significant benefit in terms of OS. Younger patients (age < median = 65 yo) have also significantly longer OS. Male and black african-american patients have significantly shorter OS. Among the treatments, only ASCT is significantly associated with better OS.

**Supp Fig. 5.**
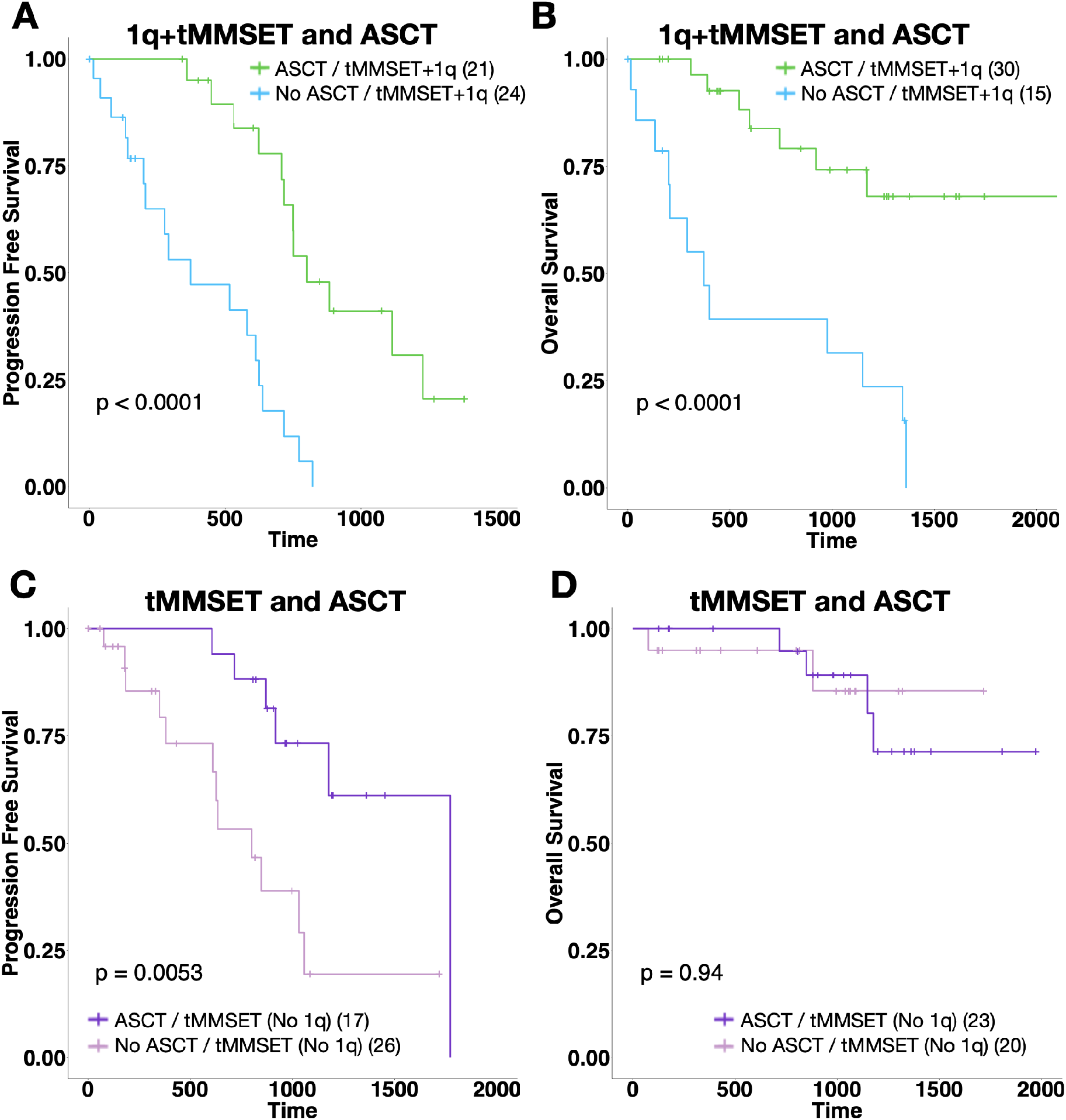
Multivariate cox-regression analysis of overall survival. **A, B.** Survival plots show that patients with 1q gain and tMMSET who received ASCT have significantly better PFS and OS compared to patients that didn’t receive ASCT. **C, D**. Survival plots show that patients with 1q gain and tMMSET who received ASCT have significantly better PFS but no significant difference in OS.

**Supp Fig. 6.**
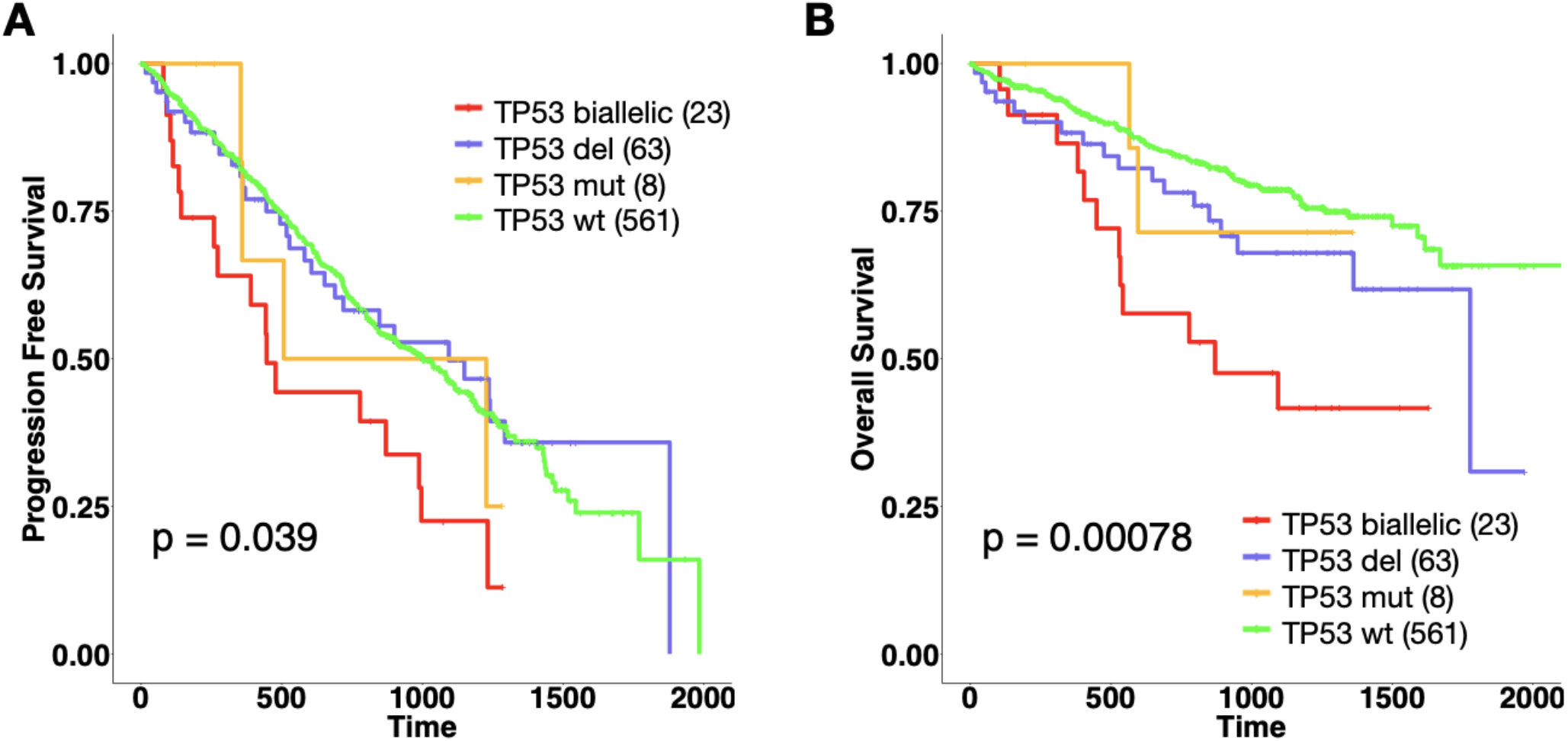
Biallelic inactivation of TP53. The plots show that biallelic inactivation of TP53, i.e. both deletion and mutation, is significantly associated with worse PFS and OS.

**Supp Fig. 7.**
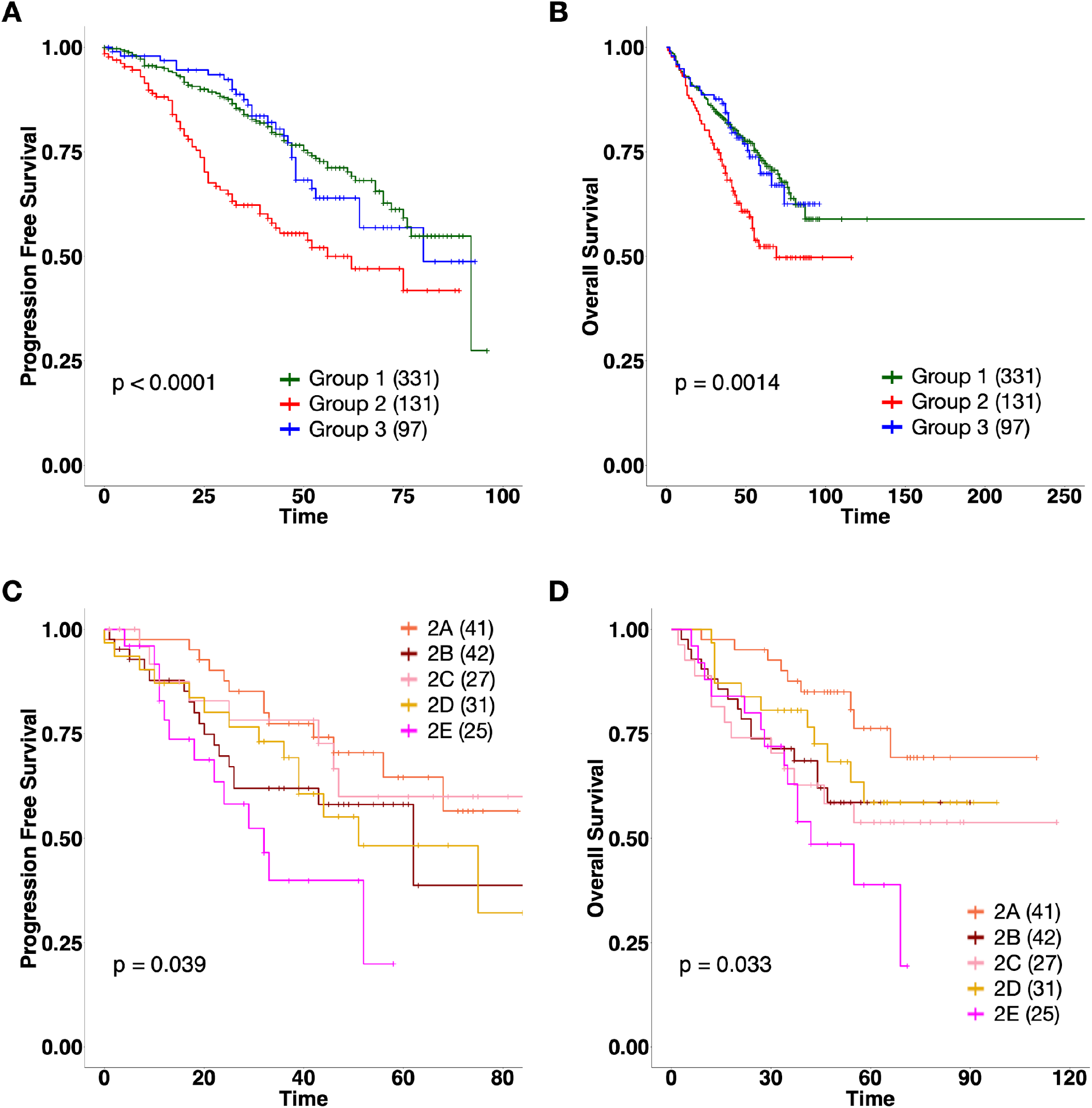
Validation of MM-PSN sub-groups in an independent dataset. A gene expression classifier was generated based on the 12 MM-PSN sub-groups and used to predict sub-groups in a cohort of newly diagnosed MM patients pre-TT2 and pre-TT3 treatment. **A, B.** Survival plots of the predicted three main groups show significantly worse PFS and OS of group 2, concordantly with MM-PSN findings. **C, D.** Survival plots of the predicted sub-groups in group 2 indicate worse PFS and OS of sub-group 2e, concordantly with MM-PSN findings.

## REFERENCES

1. Kumar SK, Rajkumar V, Kyle RA, et al. Multiple myeloma. Nat Rev Dis Primers. 2017;3:17046.

2. American Cancer Society. Cancer Facts and Figures 2019. 2019.

3. Bazarbachi AH, Al Hamed R, Malard F, Harousseau J-L, Mohty M. Relapsed refractory multiple myeloma: a comprehensive overview. Leukemia. 2019;33(10):2343–2357.

4. Multiple myeloma: 2018 update on diagnosis, risk-stratification, and management. Am. J. Hematol. 2018;93(8):981–1114.

5. Morgan GJ, Walker BA, Davies FE. The genetic architecture of multiple myeloma. Nat. Rev. Cancer. 2012;12(5):335–348.

6. Lohr JG, Stojanov P, Carter SL, et al. Widespread genetic heterogeneity in multiple myeloma: implications for targeted therapy. Cancer Cell. 2014;25(1):91–101.

7. Bolli N, Avet-Loiseau H, Wedge DC, et al. Heterogeneity of genomic evolution and mutational profiles in multiple myeloma. Nat. Commun. 2014;5:2997.

8. Laganà A, Beno I, Melnekoff D, et al. Precision Medicine for Relapsed Multiple Myeloma on the Basis of an Integrative Multiomics Approach. JCO Precis Oncol. 2018;2018.:

9. Bergsagel PL, Kuehl WM. Molecular pathogenesis and a consequent classification of multiple myeloma. J. Clin. Oncol. 2005;23(26):6333–6338.

10. Zhan F, Huang Y, Colla S, et al. The molecular classification of multiple myeloma. Blood. 2006;108(6):2020–2028.

11. Broyl A, Hose D, Lokhorst H, et al. Gene expression profiling for molecular classification of multiple myeloma in newly diagnosed patients. Blood. 2010;116(14):2543–2553.

12. Laganà A, Perumal D, Melnekoff D, et al. Integrative network analysis identifies novel drivers of pathogenesis and progression in newly diagnosed multiple myeloma. Leukemia. 2018;32(1):120–130.

13. Bolli N, Biancon G, Moarii M, et al. Analysis of the genomic landscape of multiple myeloma highlights novel prognostic markers and disease subgroups. Leukemia. 2017;

14. Pai S, Bader GD. Patient Similarity Networks for Precision Medicine. J. Mol. Biol. 2018;430(18 Pt A):2924–2938.

15. Cavalli FMG, Remke M, Rampasek L, et al. Intertumoral Heterogeneity within Medulloblastoma Subgroups. Cancer Cell. 2017;31(6):737–754.e6.

16. Cancer Genome Atlas Research Network. Integrated Genomic Characterization of Pancreatic Ductal Adenocarcinoma. Cancer Cell. 2017;32(2):185–203.e13.

17. Wang B, Mezlini AM, Demir F, et al. Similarity network fusion for aggregating data types on a genomic scale. Nat. Methods. 2014;11(3):333–337.

18. Pitroda SP, Khodarev NN, Huang L, et al. Integrated molecular subtyping defines a curable oligometastatic state in colorectal liver metastasis. Nat. Commun. 2018;9(1):1793.

19. Shen R, Mo Q, Schultz N, et al. Integrative subtype discovery in glioblastoma using iCluster. PLoS One. 2012;7(4):e35236.

20. Mo Q, Shen R, Guo C, et al. A fully Bayesian latent variable model for integrative clustering analysis of multi-type omics data. Biostatistics. 2018;19(1):71–86.

21. Robinson MD, McCarthy DJ, Smyth GK. edgeR: a Bioconductor package for differential expression analysis of digital gene expression data. Bioinformatics. 2010;26(1):139–140.

22. Reimand J, Arak T, Adler P, et al. g:Profiler—a web server for functional interpretation of gene lists (2016 update). Nucleic Acids Res. 2016;44(W1):W83–W89.

23. Merico D, Isserlin R, Stueker O, Emili A, Bader GD. Enrichment map: a network-based method for gene-set enrichment visualization and interpretation. PLoS One. 2010;5(11):e13984.

24. Su G, Morris JH, Demchak B, Bader GD. Biological network exploration with Cytoscape 3. Curr. Protoc. Bioinformatics. 2014;47:8.13.1–24.

25. Fabregat A, Jupe S, Matthews L, et al. The Reactome Pathway Knowledgebase. Nucleic Acids Res. 2018;46(D1):D649–D655.

26. Tsherniak A, Vazquez F, Montgomery PG, et al. Defining a Cancer Dependency Map. Cell. 2017;170(3):564–576.e16.

27. Meyers RM, Bryan JG, McFarland JM, et al. Computational correction of copy number effect improves specificity of CRISPR-Cas9 essentiality screens in cancer cells. Nat. Genet. 2017;49(12):1779–1784.

28. Schmidt TM, Barwick BG, Joseph N, et al. Gain of Chromosome 1q is associated with early progression in multiple myeloma patients treated with lenalidomide, bortezomib, and dexamethasone. Blood Cancer J. 2019;9(12):94.

29. Shah V, Sherborne AL, Walker BA, et al. Prediction of outcome in newly diagnosed myeloma: a meta-analysis of the molecular profiles of 1905 trial patients. Leukemia. 2018;32(1):102–110.

30. Rajkumar SV, Vincent Rajkumar S. Multiple myeloma: 2020 update on diagnosis, risk-stratification and management. American Journal of Hematology. 2020;95(5):548–567.

31. Palumbo A, Avet-Loiseau H, Oliva S, et al. Revised International Staging System for Multiple Myeloma: A Report From International Myeloma Working Group. J. Clin. Oncol. 2015;33(26):2863–2869.

32. Greipp PR, Miguel JS, Durie BGM, et al. International staging system for multiple myeloma. J. Clin. Oncol. 2005;23(15):3412–3420.

33. Chng WJ, Gualberto A, Fonseca R. IGF-1R is overexpressed in poor-prognostic subtypes of multiple myeloma. Leukemia. 2006;20(1):174–176.

34. Bataille R, Robillard N, Avet-Loiseau H, Harousseau J-L, Moreau P. CD221 (IGF-1R) is aberrantly expressed in multiple myeloma, in relation to disease severity. Haematologica. 2005;90(5):706–707.

35. Laddha SV, da Silva EM, Robzyk K, et al. Integrative Genomic Characterization Identifies Molecular Subtypes of Lung Carcinoids. Cancer Res. 2019;79(17):4339–4347.

36. Mo Q, Wang S, Seshan VE, et al. Pattern discovery and cancer gene identification in integrated cancer genomic data. Proc. Natl. Acad. Sci. U. S. A. 2013;110(11):4245–4250.

37. Madhukar NS, Khade PK, Huang L, et al. A Bayesian machine learning approach for drug target identification using diverse data types. Nat. Commun. 2019;10(1):5221.

38. Pai S, Hui S, Isserlin R, et al. netDx: interpretable patient classification using integrated patient similarity networks. Mol. Syst. Biol. 2019;15(3):e8497.

39. Parimbelli E, Marini S, Sacchi L, Bellazzi R. Patient similarity for precision medicine: A systematic review. J. Biomed. Inform. 2018;83:87–96.

40. Muthuswami M, Ramesh V, Banerjee S, et al. Breast tumors with elevated expression of 1q candidate genes confer poor clinical outcome and sensitivity to Ras/PI3K inhibition. PLoS One. 2013;8(10):e77553.

41. Chen L, Chan THM, Guan X-Y. Chromosome 1q21 amplification and oncogenes in hepatocellular carcinoma. Acta Pharmacol. Sin. 2010;31(9):1165–1171.

42. Najfeld V, Tripodi J, Scalise A, et al. Jumping translocations of the long arms of chromosome 1 in myeloid malignancies is associated with a high risk of transformation to acute myeloid leukaemia. Br. J. Haematol. 2010;151(3):288–291.

43. Marcellino BK, Hoffman R, Tripodi J, et al. Advanced forms of MPNs are accompanied by chromosomal abnormalities that lead to dysregulation of TP53. Blood Adv. 2018;2(24):3581–3589.

44. Mohan M, Weinhold N, Schinke C, et al. Daratumumab in high-risk relapsed/refractory multiple myeloma patients: adverse effect of chromosome 1q21 gain/amplification and GEP70 status on outcome. Br. J. Haematol. 2020;189(1):67–71.

45. Walker BA, Boyle EM, Wardell CP, et al. Mutational Spectrum, Copy Number Changes, and Outcome: Results of a Sequencing Study of Patients With Newly Diagnosed Myeloma. J. Clin. Oncol. 2015;33(33):3911–3920.

46. Walker BA, Mavrommatis K, Wardell CP, et al. A high-risk, Double-Hit, group of newly diagnosed myeloma identified by genomic analysis. Leukemia. 2019;33(1):159–170.

47. Chng WJ, on behalf of the International Myeloma Working Group, Dispenzieri A, et al. IMWG consensus on risk stratification in multiple myeloma. Leukemia. 2014;28(2):269–277.

48. Tagde A, Rajabi H, Bouillez A, et al. MUC1-C drives MYC in multiple myeloma. Blood. 2016;127(21):2587–2597.

49. Kawano T, Ahmad R, Nogi H, et al. MUC1 oncoprotein promotes growth and survival of human multiple myeloma cells. Int. J. Oncol. 2008;33(1):153–159.

50. Yin L, Tagde A, Gali R, et al. MUC1-C is a target in lenalidomide resistant multiple myeloma. British Journal of Haematology. 2017;178(6):914–926.

51. Leshchenko VV, Kuo P-Y, Jiang Z, et al. Harnessing Noxa demethylation to overcome Bortezomib resistance in mantle cell lymphoma. Oncotarget. 2015;6(29):27332–27342.

52. Tahir SK, Smith ML, Hessler P, et al. Potential mechanisms of resistance to venetoclax and strategies to circumvent it. BMC Cancer. 2017;17(1):399.

53. Jin S, Cojocari D, Purkal JJ, et al. 5-Azacitidine Induces NOXA to Prime AML Cells for Venetoclax-Mediated Apoptosis. Clin. Cancer Res. 2020;

54. Liu Y, Mondello P, Erazo T, et al. NOXA genetic amplification or pharmacologic induction primes lymphoma cells to BCL2 inhibitor-induced cell death. Proc. Natl. Acad. Sci. U. S. A. 2018;115(47):12034–12039.

55. Kumar S, Kaufman JL, Gasparetto C, et al. Efficacy of venetoclax as targeted therapy for relapsed/refractory t (11; 14) multiple myeloma. Blood, The Journal of the American Society of Hematology. 2017;130(22):2401–2409.

