## Supplementary Methods for "Patient Similarity Network of Multiple Myeloma Identifies Patient Sub-groups with Distinct Genetic and Clinical Features"

### SM1. Datasets acquisition and primary data generation

Whole-Genome Seq (WGS), Whole-Exome Seq (WES) and RNA-Seq data for 655 patients enrolled in the MMRF CoMMpass study was provided by MMRF and is available on the dbGaP database (http://www.ncbi.nlm.nih.gov/gap) under accession number phs000748. All RNA and DNA samples were isolated at the North American Biobank (VARI, Grand Rapids, Michigan) and provided to the genomic characterization center at the Translational Genomics Research Institute (TGen), Phoenix, Arizona. All libraries were sequenced on Illumina HiSeq2000 or HiSeq2500 using Illumina HiSeq v3 or v4 chemistry. Whole-Exome and RNA assays were sequenced using paired-end 83x83bp (TruSeqExome/RNA and Agilent V5+UTR single-plex) or 82x82bp (Agilent V5+UTR 8-plex pools) sequencing. Whole-Genome Long-insert assays were sequenced using 86x86bp sequencing. mRNA unstranded libraries were created from tumor RNA only (CD138+ cells). Whole-Exome and Whole-Genome libraries were created from matched tumor (CD138+ cells) and peripheral blood samples.

All primary data were generated at TGen as follows. Fastq files were aligned to the GRCh37 reference human genome and all annotation and gene models were based on Ensembl version 74. Whole-Exome and Whole-Genome fastq files were aligned to the reference genome using BWA (0.7.8)[1]. Sequence alignment map (SAM) files were converted to binary alignment maps (BAM) files using Samtools (0.1.19)[2]. Lane level quality recalibration was performed using PICARD (1.111). Duplicate fragments were identified and marked using

PICARD (1.111). Joint-Indel realignment was performed on the matched tumor and constitutional bam files using GATK (GenomeAnalysisTK-3.1-1)[3].

RNAseq fastq files were aligned to the reference genome using STAR (2.3.1z)[4]. Gene expression estimates were calculated using HT-Seq (0.6.0) counts[5].

Somatic single nucleotide variants (SNVs) and small INDELs were identified from matched tumor-normal WES data using an integrated approach leveraging three different somatic variant callers: Seurat v2.6, Strelka v1.0.13, and MuTect v1.1.4[6–8]. Somatic mutations were filtered to include calls made by at least two of the three callers.

Copy number alterations (CNAs) were identified using a CoMMpass-specific version of TGen's in-house copy number tool tCoNuT_COMMPASS. Relative copy number was determined as the log2 difference between the normal and tumor normalized coverage, where normalization is the mean coverage across a 2 kb window divided by the genome-wide mode coverage. We additionally used the tool Gistic 2.023 to identify regions of the genome that were significantly amplified or deleted across multiple samples. CNAs were classified as broad (arm-level) or focal (band/sub-band level)[9].

Structural variants were identified from the whole-genome long-insert assay using the tools DELLY and an internally developed TGen caller.

To detect gene fusion transcripts RNA-Seq data was aligned using TopHat2 v2.0.11[10]. TopHat-Fusion post was set to report fusions[11]. Potential fusions called by TopHat-Fusion were subject to a guided-fusion assembly approach to assemble each of the potential fusion transcripts using the tool Trinity 2.2 for de novo assembly of RNA-Seq data and bwa-mem version 0.7.8 for alignment to the fusion reference[12]. Potential fusions were also independently validated in the long-insert WGS data by traversing through a series of windows initially set to a size of 1000 base pairs.

**Generation and clustering of Patient Similarity Network**

MM-PSN was generated using the SNF method implemented in the R package SNFtool (v2.3.0)[13]. The tool was run on 655 MM tumor samples using gene expression (50,495 genes), SNV (57,736 mutations), gene fusion (13,682 fusions), focal CNA (93 features) and Broad CNA data (39 features), using the parameters K = 50, alpha = 0.6, T = 50. The fused matrix obtained from the SNF function was clustered using the Spectral Clustering method implemented in the SNFtool package, setting k=2 to 15 (k = number of clusters). We selected three as the optimal number of clusters which maximized the Eigen Gap and minimized the Rotation Cost, as suggested by the authors of SNF (**Fig. 1D**). For each of these three groups, we ran SNFtool again independently with the following parameters:

Group 1: K = 40; alpha = 0.6; T = 50

Group 2: K = 60; alpha = 0.6; T = 50

Group 3: K = 20; alpha = 0.6; T = 50

We then employed spectral clustering on the fused similarity matrices, setting k=2 to 8, and selected four, five and three as the optimal number of sub-groups for groups 1, 2 and 3, respectively.

**Network Visualization**

We retrieved all the patient pairs from the fused similarity matrix W returned by SNF and retained only those with similarity greater than the third quartile values of all possible pairs, for improved visualization. These filtered pairs were then imported in Cytoscape (v3.7.2)[14]. We used the edge-weighted spring embedded layout for visualizing the edges shown in **Fig. 1A**.

**Differential feature analysis and functional enrichment**

We calculated the differentially expressed (DE) genes for the 12 subgroups by comparing their expression in each sub-group with the average expression in the other 11 subgroups using the R package edgeR[15]. For pathway analysis, we retained the upregulated genes in each subgroup (logFC >= 1.5 and FDR <0.05) and excluded the immunoglobulin genes. Pathway enrichment analysis was performed using the tool g:Profiler and visualized as Enrichment Map in Cytoscape[16,17] . Pathway definitions were retrieved from Reactome[18]. The CERES scores estimating gene-dependency levels from CRISPR-Cas9 essentiality screens were retrieved from DepMap[19,20]. We considered as essential genes with CERES score < -0.5, as suggested in the DepMap portal.

**MM-PSN classifier**

Data from the 655 samples was divided into training and validation sets in the ratio of 70:30. The classifier was built by combining three types of features, i.e. actual copy number of focal chromosome bands, translocation calls and gene expression. First, feature selection was performed on copy number values using a Recursive Feature Elimination (RFE) strategy. XGBoost was used as an external estimator that assigned weights to the features. First, we filtered out highly correlated focal bands (correlation > 0.95) and trained XGBoost on the remaining 85 bands. The importance of each feature was measured in terms of the weights assigned by the classifier. Then, the five least important features with minimum weight were pruned from the current set of features. This process was recursively reiterated on the trimmed set until the required number of features was eventually reached. We selected the top 50 features which maximized the performance in terms of weighted recall. Then, the selected features were converted into categorical variables according to the following criteria: Between 0 and 0.5: Loss; Between 0.5 and 1.5: Deletion; Between 1.5 and 2.5: Normal; Between 2.5 and 3.5: Gain; >3.5: Amplification. The features were then further one-Hot encoded to be fed into the classifier.

For gene expression, we first selected the top 100 genes with the highest NMI (Normalized mutual information) for each of the three main groups and their subgroups. The VST (Variance Stabilizing Transformation) normalized gene expression features of the training dataset were scaled to z-scores[21]. The validation gene expression dataset was normalized using the mean and standard deviation of training data. Then, we used a two-step feature selection method: first, the features were filtered based on a linear support vector machine with L1 norm (SVC-L1), then an RFE approach using XGBoost was applied similarly to as described above for CNA data. Finally, a total of 109 gene expression features were selected and converted to categorical variables as per the following criteria: Z-score >1.5: Upregulated; Z score < -1.5: Downregulated; Z-Score >-1.5 and Zscore <1.5: Normal expression. The categorical expression features were then OneHot encoded.

We then combined the one hot encoded CNA, Expression and all the 8 translocation calls side by side and used a stacking classifier with Random forest, Xgboost and linear SVC as the base learners. Then, a linear support vector machine was used as a final estimator to compute the final prediction of sub-groups taking the input from the three base classifiers. The performance on training data was calculated using 3-fold cross validation. The final model was developed using full training data and tested on 30% validation dataset separated upfront from the full dataset. Fig. SM1 and SM2 show the Precision vs Recall curves for the training and test sets, respectively.

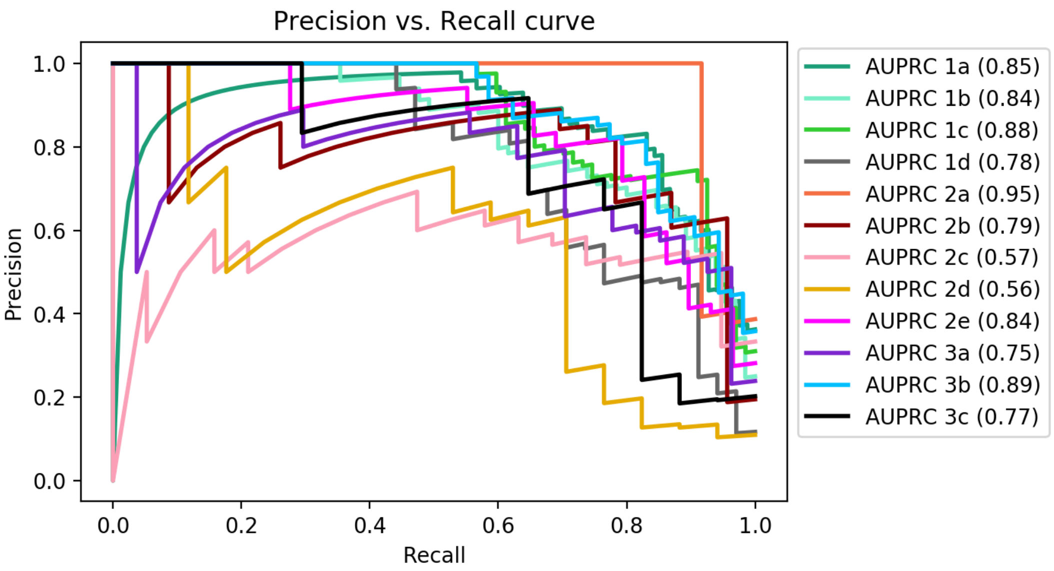


**Fig. SM1**. Precision vs Recall curve for the training set.

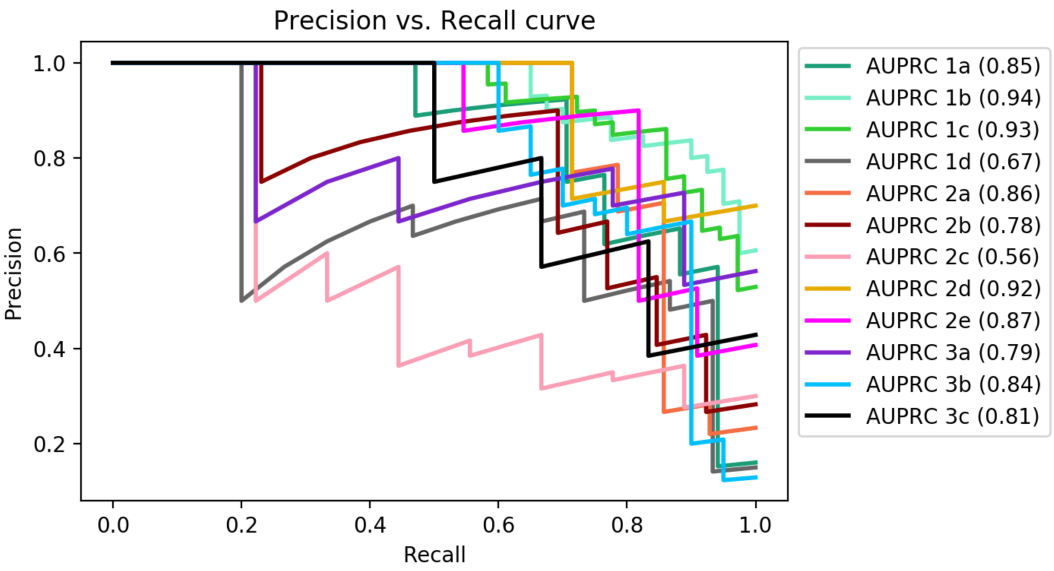


**Fig. SM2**. Precision vs Recall curve for the test set.

**Network validation**

To validate MM-PSN groups and sub-groups on an independent gene expression dataset, we trained classification models on the 655 samples using gene expression data only. The 12 sub-groups obtained by spectral clustering on the SNF fused similarity matrix were taken as the ground truth for label assignment. We implemented Logistic Regression, Random Forest and Support Vector Machines (SVM) and compared their performances. The best models were selected using 10-fold cross validation at the group level and 5-fold cross validation at the sub-group level on the training data. The validation data was generated with Affymetrix GeneChip U133 plus 2.0 arrays from Shaughnessy et al and was retrieved from NCBI GEO (GSE2658).

**REFERENCES**


1. Li H, Durbin R. Fast and accurate short read alignment with Burrows–Wheeler transform. Bioinformatics. 2009;25(14):1754–1760.
2. Li H, Handsaker B, Wysoker A, et al. The Sequence Alignment/Map format and SAMtools. Bioinformatics. 2009;25(16):2078–2079.
3. Van der Auwera GA, Carneiro MO, Hartl C, et al. From FastQ data to high-confidence variant calls: the genome analysis toolkit best practices pipeline. Curr. Protoc. Bioinformatics. 2013;43(1):11–10.
4. Dobin A, Davis CA, Schlesinger F, et al. STAR: ultrafast universal RNA-seq aligner. Bioinformatics. 2013;29(1):15–21.
5. Anders S, Pyl PT, Huber W. HTSeq--a Python framework to work with high-throughput sequencing data. Bioinformatics. 2015;31(2):166–169.
6. Christoforides A, Carpten JD, Weiss GJ, et al. Identification of somatic mutations in cancer through Bayesian-based analysis of sequenced genome pairs. BMC Genomics. 2013;14:302.
7. Saunders CT, Wong WSW, Swamy S, et al. Strelka: accurate somatic small-variant calling from sequenced tumor–normal sample pairs. Bioinformatics. 2012;28(14):1811–1817.
8. Cibulskis K, Lawrence MS, Carter SL, et al. Sensitive detection of somatic point mutations in impure and heterogeneous cancer samples. Nat. Biotechnol. 2013;31(3):213–219.
9. Mermel CH, Schumacher SE, Hill B, et al. GISTIC2.0 facilitates sensitive and confident localization of the targets of focal somatic copy-number alteration in human cancers. Genome Biol. 2011;12(4):R41.
10. Trapnell C, Pachter L, Salzberg SL. TopHat: discovering splice junctions with RNA-Seq. Bioinformatics. 2009;25(9):1105–1111.
11. Kim D, Salzberg SL. TopHat-Fusion: an algorithm for discovery of novel fusion transcripts. Genome Biol. 2011;12(8):R72.
12. Haas BJ, Papanicolaou A, Yassour M, et al. De novo transcript sequence reconstruction from RNA-seq using the Trinity platform for reference generation and analysis. Nat. Protoc. 2013;8(8):1494–1512.
13. Wang B, Mezlini AM, Demir F, et al. Similarity network fusion for aggregating data types on a genomic scale. Nat. Methods. 2014;11(3):333–337.
14. Su G, Morris JH, Demchak B, Bader GD. Biological network exploration with Cytoscape 3. Curr. Protoc. Bioinformatics. 2014;47:8.13.1–24.
15. Robinson MD, McCarthy DJ, Smyth GK. edgeR: a Bioconductor package for differential expression analysis of digital gene expression data. Bioinformatics. 2010;26(1):139–140.
16. Reimand J, Kolde R, Arak T. gProfileR: Interface to the'g: Profiler'Toolkit. R package version 0. 6. 2018;7.:
17. Merico D, Isserlin R, Stueker O, Emili A, Bader GD. Enrichment map: a network-based method for gene-

set enrichment visualization and interpretation. PLoS One. 2010;5(11):e13984.

18. Fabregat A, Jupe S, Matthews L, et al. The Reactome Pathway Knowledgebase. Nucleic Acids Res. 2018;46(D1):D649–D655.

19. Tsherniak A, Vazquez F, Montgomery PG, et al. Defining a Cancer Dependency Map. Cell. 2017;170(3):564–576.e16.

20. Meyers RM, Bryan JG, McFarland JM, et al. Computational correction of copy number effect improves specificity of CRISPR-Cas9 essentiality screens in cancer cells. Nat. Genet. 2017;49(12):1779–1784.

21. Anders S, Huber W. Differential expression of RNA-Seq data at the gene level--the DESeq package. Heidelberg, Germany: European Molecular Biology Laboratory (EMBL). 2012;10:f1000research.
